# Anti-EGFR aptamer exhibits direct anti-cancer effects in NSCLC cells harboring EGFR L858R mutations

**DOI:** 10.1101/2024.04.01.587576

**Authors:** Brian J. Thomas, Sania Z. Awan, Trupti Joshi, Mark A. Daniels, David Porciani, Donald H. Burke

## Abstract

Non-small cell lung cancer (NSCLC) adenocarcinoma (LUAD) is a leading cause of death worldwide. Activating mutations in the tyrosine kinase domain of the oncogene epidermal growth factor receptor (EGFR) are responsible for ∼10-50% of all LUAD cases. Although EGFR tyrosine kinase inhibitors (TKIs) have been effective in prolonging NSCLC patient survival and quality of life, acquired resistance mechanisms and disease progression are inevitable. Contemporary second- and third-line treatments, such as immunotherapy, remain ineffective for these patients, presenting a clear and unmet need for alternative or adjuvant therapeutics for the treatment of mutant EGFR positive NSCLC. Here we show that an anti-EGFR aptamer (*EGFRapt*) decreases viability of NSCLC cell lines harboring the L858R ± T790M mutation in EGFR but not cell lines harboring wild-type or exon 19 deletions. In a humanized xenograft mouse model of NSCLC, *EGFRapt* decreased tumor burden compared to controls when delivered intratumorally over multiple doses. To elucidate the mechanism by which *EGFRapt* exerts these effects, we monitored cellular processes associated with kinase-dependent and kinase-independent mechanisms and found that the anti-cancer effects of *EGFRapt* are cell line dependent, inhibiting cellular proliferation in one cell line and inducing cell death in another. Post hoc transcriptomics analysis supported these findings and provided additional mechanistic insights. Overall, these data establish that *EGFRapt* has direct anti-cancer activity in mutant EGFR positive NSCLC via targetable mechanisms that are independent of existing approaches, and they provide a foundation for further development of nucleic acid-based therapies that target EGFR.

## Introduction

As a leading cause of death in the United States and across the globe, lung cancer will kill more people in 2024 than breast, prostate, and colon cancer combined^1,2^. Lung adenocarcinoma (LUAD) makes up nearly 40% of all NSCLC diagnoses and is typically defined by genetic alterations in oncogenes such as *EGFR, KRAS,* and *ALK*, among others, that lead to dysregulated cellular metabolism and sustained proliferative signaling^3,4^. Primary mutations in the tyrosine kinase domain (TKD) of epidermal growth factor receptor (EGFR) account for ∼10-50% of LUAD cases, depending on race and ethnicity^5–7^. TKD Exon 21 L858R mutations (40%), exon 19 deletions (Ex19del, 45%), and exon 20 insertions (Ex20ins, 5-10%), which collectively make up the majority of mutations seen clinically^6,8^, lead to protein dimerization and constitutive activation of signaling even in the absence of canonical ligands^9^. Such ‘inside-out’ receptor dimerization has been shown to increase canonical, kinase-dependent signaling through EGFR, promoting cancer cell growth and proliferation. However, more recent reports have described non-canonical or kinase-independent functions of wild-type and mutant EGFR that instead promote cancer cell survival^10,11^. These non-canonical functions are typically a result of metabolic reprogramming (*e.g.*, stabilization of glucose transporters) or receptor trafficking to other cellular compartments such as ER, mitochondria, or the nucleus^11–13^. Mutations in the TKD of EGFR have been reported to cause altered protein interactions (*e.g.*, heterodimerization) and aberrant trafficking, affecting both canonical and non-canonical functions and negatively impacting patient survival^14–17^. Nonetheless, the many functions of mutant EGFR in cancer progression suggests that opportunities exist for alternative or novel therapeutic intervention.

Despite effective EGFR targeted therapies such as small molecule inhibitors (*e.g.*, Osimertinib, a tyrosine kinase inhibitor or TKI) and antibodies (*e.g.*, amivantamab, an EGFR/c-Met bispecific antibody), NSCLC patients treated with these reagents commonly relapse within 12-18 months^18,19^. Acquired therapeutic resistance and progressive disease are commonly due to the inherent heterogeneity of tumor cells and the tumor microenvironment (TME)^20,21^. Resistance mechanisms most common to anti-EGFR therapy include (i) secondary/tertiary mutations in the TKD that disrupt TKI binding affinity (*e.g.*, EGFR T790M or C797X mutations), (ii) downregulation of the targeted oncogene, (iii) clonal selection of cells with mutation or amplification in an untargeted oncogene (*e.g.*, activation of downstream oncogenes such as KRAS or other oncogenic pathways such as c-Met and AXL), or (iv) phenotype transformation (*e.g.*, epithelial mesenchymal transition or histological transformation to small cell lung cancer)^22–25^. The five-year survival for relapsed patients remains low, especially as immunotherapy and salvage chemotherapy, which are the go-to second and third-line treatment options for TKI resistant NSCLC, have proven to be ineffective^24,26,27^ or to induce life-threatening side effects^28^. This again highlights an unmet need for the development of alternative or adjuvant NSCLC therapies.

The federal Cancer Moonshot^29,30^ and the National Cancer Institute’s (NCI) National Cancer Plan (nationalcancerplan.cancer.gov) initiatives promote a goal of greater than fifty percent reduction in cancer mortality by the year 2050. To achieve these goals, the development of novel therapeutics for rare, difficult to treat, and resistant cancers is imperative. Specifically, the development of effective treatments for these cancers should (i) harbor unique mechanisms of actions, (ii) have low toxicity (*i.e.*, produce minimal side effects), and (iii) be accessible to and obtainable by all people (*i.e.*, minimize healthcare barriers). Given the continued improvements in our understanding in the use of oligonucleotides as clinical reagents^31–34^, aptamer technology^35–37^ has the potential to play a role in this novel therapeutic developmental pipeline.

Aptamers are short strands of oligonucleotides (DNA or RNA) that fold into a sequence-specific three-dimensional conformation, enabling them to recognize their molecular targets with high specificity and affinity (Kd typically nM to high pM). Aptamers are typically discovered through a process called systematic evolution of ligands through exponential enrichment (SELEX), where large oligonucleotide pools are narrowed down through iterative incubations with proteins, cell lines, or animal models of interest, with each step enriching the population for species with appropriate molecular recognition characteristics. As aptamers (∼10-20 kDa) are typically smaller than monoclonal antibodies (∼150 kDa) or antibody fragments (Fab; ∼50 kDa), they have the ability to bind unique epitopes and potentially provide an alternative anti-cancer mechanism to current therapies. Aptamers have also been reported to induce minimal toxicity in humans (as they are not recognized by the adaptive immune system), are relatively inexpensive to produce, and are readily scalable with low batch-to-batch variation^35,38^. Each of these properties lend well to the three goals of novel therapeutic development mentioned above. However, aptamers do harbor pharmacokinetic limitations that have precluded their translation to the clinic for cancer therapy. To address these bottlenecks, our lab and others are currently screening modified aptamers^37,39–41^ and improving selection technology (*e.g.*, enzymes used in canonical SELEX)^42^ to engineer or develop aptamers with better pharmacokinetic properties.

Multiple well-characterized aptamers have been selected toward targets that are upregulated on NSCLC LUAD, including EGFR, and these have been used to disrupt biology and to target delivery of anti-cancer reagents^43–45^. We and others have shown that the 2’-fluoropryrimidine (2’FY) anti-EGFR aptamer, MinE07^43,46^ (herein referred to as *EGFRapt*), is a reliable aptamer that binds the extracellular domain III of EGFR and targets both WT and mutant EGFR NSCLC *in vitro* and *in vivo*^47–49^. We have recently utilized *EGFRapt* in the development of barcoded aptamer technology (BApT), which allowed multiplexed *in vivo* screening of pre-defined aptamer formulations to identify superior targeting reagents^50^. As an established tumor targeting reagent, *EGFRapt* can be confidently developed to facilitate its therapeutic use in EGFR positive cancers, either as a direct acting anti-cancer reagent or as a therapeutic cargo carrier and delivery reagent.

In this work, we show that *EGFRapt* has direct acting anti-cancer effects in NSCLC LUAD cell lines. Specifically, we screened multiple LUAD cells lines harboring either WT or mutant EGFR and found that *EGFRapt* decreases cell viability in two cell lines (H3255 and H1975) that harbor the oncogenic EGFR L858R ± T790M TKD mutations. We show that this anti-cancer effect is retained *in vivo* in an H1975-derived subcutaneous xenograft model of NSCLC when *EGFRapt* is injected locally. Using both benchtop and computational methods, we show that the mechanism by which *EGFRapt* induces this anti-cancer effect differs between the two cell lines and is distinct from the current clinically relevant anti-EGFR therapeutic toolbox. Together, our findings provide insight into additional mechanisms that may be targeted in the treatment of NSCLC and establish *EGFRapt* as a potential anti-cancer reagent for further therapeutic development and testing.

## Results

### EGFRapt decreases cell viability in vitro in NSCLC cell lines harboring EGFR L858R mutations

As we have previously shown that *EGFRapt* binds to the extracellular domain III of EGFR at sites that overlap the clinically relevant monoclonal antibodies (mAb) cetuximab and nimotuzumab, we hypothesized that this aptamer might have anti-cancer activity by regulating canonical or non-canonical EGFR pathways^11^. Therefore, we evaluated the impact of *EGFRapt* on four NSCLC cell lines harboring moderate (H1975, H820) to high (A549, H3255) levels of wild-type (WT) or mutant EGFR and one breast cancer cell line (MCF7) harboring very low levels of WT EGFR. Each cell line was exposed to 3 µM *EGFRapt* or 2’FY RNA non-binding control aptamer (extended version of C36^49^; control apt) for 48 h, and then cell viability was measured by incubating with MTS cell viability reagent (**Figure 1A**). Two cell lines that each harbor EGFR L858R, either along with (H1975) or without (H3255) the T790M secondary mutation, showed a 20-30% decrease in cell viability compared to vehicle or to treatment with control apt (**Figure 1B**). In contrast, NSCLC cell lines A549 and H820, which harbor WT and Ex19 del/T790M mutations, respectively, showed no change in cell viability upon treatment with *EGFRapt* relative to treatment with control apt (**Figure 1B**). The breast cancer cell line, MCF7, exhibited increased cell viability upon treatment with either *EGFRapt* or control apt (**Figure 1B**), but with no difference between the two treatments, implying a non-specific stimulation unrelated to EGFR targeting. To further control for non-specific effects of oligonucleotides, all NSCLC cell lines were also exposed to a DNA non-binding control aptamer (scDW4; **Figure S1A-D**). Under these conditions, negligible change in cell viability was noted compared to vehicle treated cells. A dose response of *EGFRapt* in H3255 cells revealed an EC50 of ∼4.8 µM (**Figure 1C**). Interestingly, H1975 cells exhibited only partial response; therefore, a constrained (cell viability = 0.6) EC50 was calculated at ∼1.8 µM (22 µM unconstrained; **Figure 1C**). In both cell lines, control apt showed minimal effect on cell viability at nearly all concentrations tested except for the highest dose (10 µM) in H1975. To ensure the partial response exhibited by H1975 cells was specific to *EGFRapt* binding, H1975 cells were also exposed to a 2’FY RNA and a DNA ‘binding’ control aptamers that bind to transferrin receptor (extended version of Waz) and c-Met (minimized CLN3), respectively, as these antigens are highly expressed (like EGFR) on this and other cancer cell lines. In contrast to *EGFRapt*, neither binding control aptamer had a significant impact on the cell viability of H1975 (**Figure S1A**). A549 cells showed no changes in cell viability when exposed to higher doses of *EGFRapt* or control aptamer (**Figure S1E**). Representative brightfield images of the dose response at 48 h showed decreased cell numbers in H1975, whereas H3255 showed rounding of cells with minimal change in cell numbers (**Figure 1D**). As only a partial response was noted in H1975 upon a single treatment with *EGFRapt*, we sought to determine whether multiple doses would further decrease cell viability in these cell lines. Indeed, four doses of 3 µM *EGFRapt* decreased cell viability by ∼40-50% in both cell lines, in contrast to only a very slight decrease in cell viability for multiple doses of control apt (15-20%) (**Figure 1E**). Together these findings established *EGFRapt* as an anti-cancer reagent for NSCLC cell lines harboring EGFR L858R ± T790M mutations but not Ex19del or WT EGFR. Screening of additional cells lines or patient samples is warranted to establish generalizability of the mutation selectivity.

**Figure 1.**
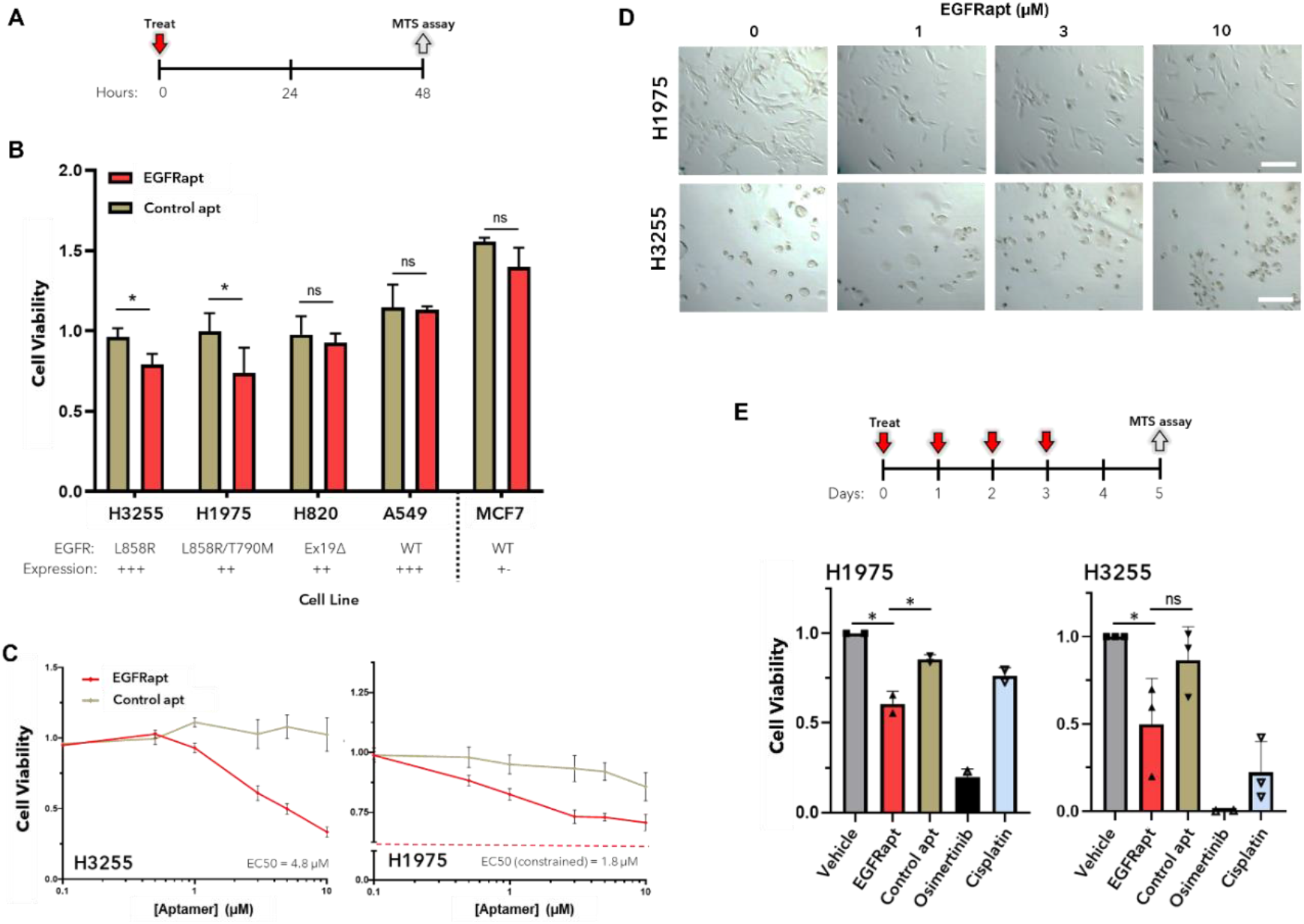
*EGFRapt* decreases cell viability in NSCLC harboring EGFR L858R mutations. (**A**) Schematic of cell viability assay using CellTiter 96® Aqueous One Solution (MTS reagent). (**B**) Indicated cell lines expressing high (H3255, A549), moderate (H1975, H820), or negligible (MCF7) levels of WT or mutant EGFR were treated with either 3 µM *EGFRapt* (red bar) or non-binding control apt (tan bar) for 48 h. Cell viability was determined using MTS reagent and plotted relative to viability of vehicle-treated cells. Two cell lines expressing L858R mutations (H3255 and H1975) exhibited decreased cell viability upon treatment with *EGFRapt* when compared to non-binding control apt. (**C**) H3255 (*left*) and H1975 (*right*) cells were treated as in B except using various doses of aptamer. H3255 cells showed a near complete loss of viability upon treatment with higher doses of *EGFRapt* (red line) compared to control apt (tan line; IC50 = 4.8 µM), whereas H1975 cells showed only a partial loss of viability (IC50 constrained = 1.8 µM; bottom at 0.6). (**D**) Bright field images of the *EGFRapt* dose response from C at 48 h. H1975 cells treated with 1, 3, or 10 µM *EGFRapt* exhibited decreased cell density compared to vehicle treatment (0 µM) whereas H3255 cells exhibited increased cell rounding with minimal changes in cell density. For B-C, relative cell viability (normalized to vehicle) is reported on the *y*-axis and plotted values represent mean ± SD for n=2-5 independent experiments, each with 2-4 technical replicates. Statistical analysis was preformed using a two tailed t-test (*p < 0.05). (**E**) H1975 and H3255 cells were subject to cell viability assay as in Figure 1B except four doses of *EGFRapt* or non-binding control apt were added in 24 h intervals (*assay schematic*). H1975 (left) or H3255 (right), were treated with four doses of 3 µM *EGFRapt* (red bar) or non-binding control apt (tan bar), or one dose of 0.1 µM Osimertinib (black bar), 2 µM Cisplatin (blue bar), or vehicle (grey bar) for 5 days, and cell viability was determined using MTS reagent. Multiple doses of *EGFRapt* further decreased cell viability in H1975 and H3255 when compared to a single dose (Figure 1B and S1). For B,C,E, relative cell viability (normalized to vehicle) is reported on the *y*-axis and plotted values represent mean ± SD for n=2-3 independent experiments, each with 2-4 technical replicates. Statistical analysis was preformed using a two tailed t-test (*p < 0.05).

### EGFRapt decreases tumor burden in vivo in NSCLC xenografts

To determine whether the anti-cancer effect observed *in vitro* was retained in a more clinically relevant cancer model, BALB/c mice harboring ectopic H1975 derived xenografts were subject to five intratumoral doses (150 pmol on days 0, 2, 4 and 200 pmol on days 6, 7) of *EGFRapt* or control apt, or a vehicle (1X DPBS), and tumor volumes were tracked until a humane endpoint was reached (**Figure 2A**). The subcutaneous tumors were soft, barely palpable, and averaged 13.3 ± 4.8 mm^3^, when the first dose was administered (day 0), approximately 12 days after engraftment. Average tumor volumes (**Figure 2B**) of individual tumor trajectories (**Figure S2A**) relative to day 0 revealed that *EGFRapt* treatment decreased tumor burden compared to control apt (p = 0.00046) or vehicle (p = 0.0007) when assessed by a simple linear regression model. The same assessment using raw tumor volumes (**Figure S2B**) also showed a significant decrease in tumor burden when compared to control apt (p = 0.0067) or vehicle (p = 0.000077). In addition to simple linear regression (**Figure S3A**), an area under the curve (AUC) model (**Figure S3B**) was also used to test tumor burden reduction in response to *EGFRapt* treatment. This analysis revealed a significant decrease only for raw tumor volumes relative to vehicle (p = 0.024) but not control apt (p = 0.17).

**Figure 2.**
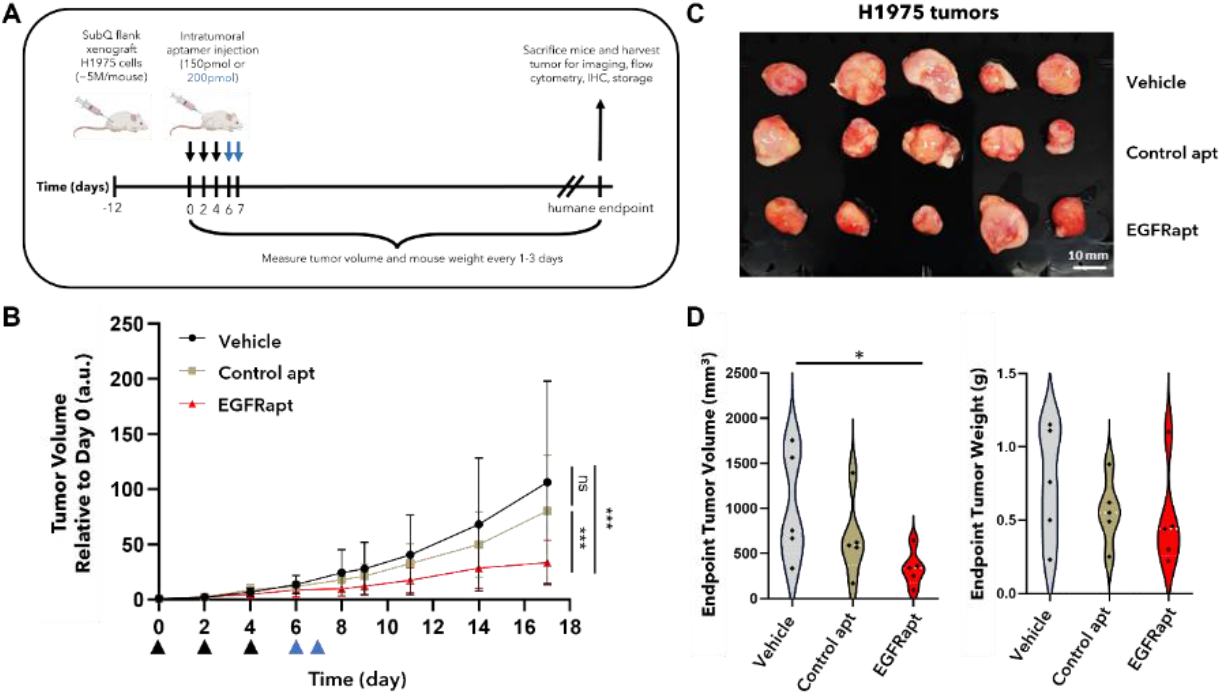
*EGFRapt* decreases tumor burden in NSCLC xenograft. (**A**) Schematic of H1975 (NSCLC) derived xenograft mouse model. (**B**) Tumor bearing mice were treated with five intratumoral doses of *EGFRapt* (red line), non-binding control apt (tan line), or vehicle (black line), and tumor burden was tracked until a humane endpoint. Tumor volume (mm^3^) relative to day 0 is plotted on the *y*-axis. Plotted values represent mean ± SD for n=5 mice from each treatment group. Tumor growth kinetics were significantly decreased in mice treated with *EGFRapt* compared to control apt or vehicle (PBS). (**C**) *Ex vivo* image of H1975 tumors from all mice at the endpoint. (**D**) *Y*-axis defines endpoint *ex vivo* tumor volumes (mm^3^; *left*) or tumor weights (grams; *right*) of mice treated with *EGFRapt* (red violin), control apt (tan violin), or vehicle (grey violin). Violin plot width represents probability and white dotted line represents mean for n=5 mice from each treatment. The majority of tumors from mice treated with *EGFRapt* were smaller than those from mice treated with control apt or vehicle. Statistical analysis was preformed using a linear regression model (B; ***p < 0.0002) or two tailed t-test (D; *p < 0.05)

At the humane endpoint (day 17), mice were sacrificed and imaged in prone and right recumbent positions (**Figure S2C**). Tumors were excised (**Figure 2C**) and *ex vivo* tumor weights and volumes were obtained (**Figure 2D**). Mice treated with *EGFRapt* exhibited significantly smaller end-point tumor volumes (mean = 338 mm^3^) when compared to vehicle (mean = 1015 mm^3^; p = 0.0465) and was less than but not significantly different from control apt (mean = 668 mm^3^; p = 0.169). End-point tumor weight was not significantly different between *EGFRapt* (mean = 0.504 g) and vehicle (mean = 0.750 g; p = 0.326) or control apt (mean = 0.558 g; p = 0.779), despite the significant difference in volume. However, the majority of the tumors (4 of 5) from *EGFRapt* treated mice weighed less than the mean tumor weight from vehicle and control apt treated mice. While not certain, the high variability in tumor size upon treatment may be due to that fact that intratumoral injection is subject to quick dispersion of injected material away from injection site (*i.e.*, tumor), especially if the injected volume is more than tumor volume, as was the case for the first two injections, where the injected volume (50 µL = 50 mm^3^) was in excess of the average tumor volume of all mice at that point (13.3 mm^3^ and 27.2 mm^3^ on day 0 and 2, respectively). Nonetheless, these findings confirmed that the anti-tumor activity of *EGFRapt* is retained in an *in vivo* model where multiple doses are delivered using local delivery methods (*i.e.*, intratumoral injection), which allow high local concentrations to be delivered to the tumor.

### Chemically modified EGFRapt retains anti-cancer effect in vitro

The 2’FY modifications of *EGFRapt* stabilize it significantly against serum nucleases (i.e., degradation). However, the current standard for nucleic acid therapeutics is that all or nearly all positions carry modifications. Superior performance could potentially emerge by introducing additional modifications in *EGFRapt* to further improve its metabolic stability. We screened two highly-modified versions of *EGFRapt*^51^ that contained a 3’ inverted dT (IdT) and forty-one (*EGFRapt* 41 mod) or forty-three (*EGFRapt* 43 mod) chemically modified positions, including twenty 2’FY and twenty-one or twenty-three 2’-*O*-methyl-purines (2’OMeR), along with six or four unmodified nucleotides, respectively (**Figure S4A**). A single 3 µM treatment with either modified *EGFRapt* showed similar anti-cancer effect compared to the original *EGFRapt* (*EGFRapt* 2’FY), decreasing viability of both H1975 and H3255 cells by ∼20% compared to vehicle, control apt (control apt 2’FY), and a fully modified version of control apt that contained 3’ IdT, seventeen 2’FY, and nineteen 2’OMeR (control apt 36 mod; **Figure S4B**). These findings, among others^51^, show that *EGFRapt* is amenable to extensive modification without loss of binding or anti-cancer activity.

### EGFRapt disrupts the cell cycle and induces apoptosis in H3255 but not in H1975

The MTS based cell viability assays above only determine whether a treatment changes the number of metabolically active cells and does not determine whether the treatment is anti-proliferative (less cell division) or cytotoxic (cell death). EGFR is known to have both a canonical (kinase-dependent) function that promotes cell growth and proliferation and a non-canonical (kinase-independent) function that promotes cell survival. Therefore, we sought to determine whether the *EGFRapt* mechanism of action was through inhibiting one or both functions.

To differentiate between these two possibilities, H3255 cells were treated with a single dose of 3 µM *EGFRapt* or control apt for 48 h cells, then fixed and stained with propidium iodide to determine the percentage of cells in the G0/G1, S, or G2/M phase of the cell cycle. *EGFRapt* treatment increased the number of cells in the S phase, similar to treatment with 1 µM 5-fluorouracil (5-FU), an S-phase specific anti-cancer therapeutic (**Figure 3A**). At the 48 h timepoint, we also analyzed the proportion of cells stained for cleaved caspase 3/7 (apoptosis) and/or Sytox (necrosis or late apoptosis). Compared to vehicle and control apt, a single dose of *EGFRapt* increased the number of cells stained with cleaved caspase 3/7 (**Figure 3B**) but not Sytox, suggesting that *EGFRapt* induces apoptosis in NSCLC cell line H3255, although it remains to be determined whether cell cycle changes are directly related to the observed increase in cell death. In contrast to these finding in H3255, cell cycle (**Figure S5A**) and cytotoxicity (**Figure S5B**) analysis in H1975 treated with *EGFRapt* revealed no difference from vehicle or binding control apt (extended version of aptamer E3^44^). Unexpectedly, the non-binding control apt significantly decreased the number of cells in G0-G1 and increased the number of cells in S phase, despite having no effect on cell viability or cytotoxicity. Nevertheless, these findings suggest that *EGFRapt* is cytotoxic and likely alters the non-canonical functions of EGFR in H3255.

**Figure 3.**
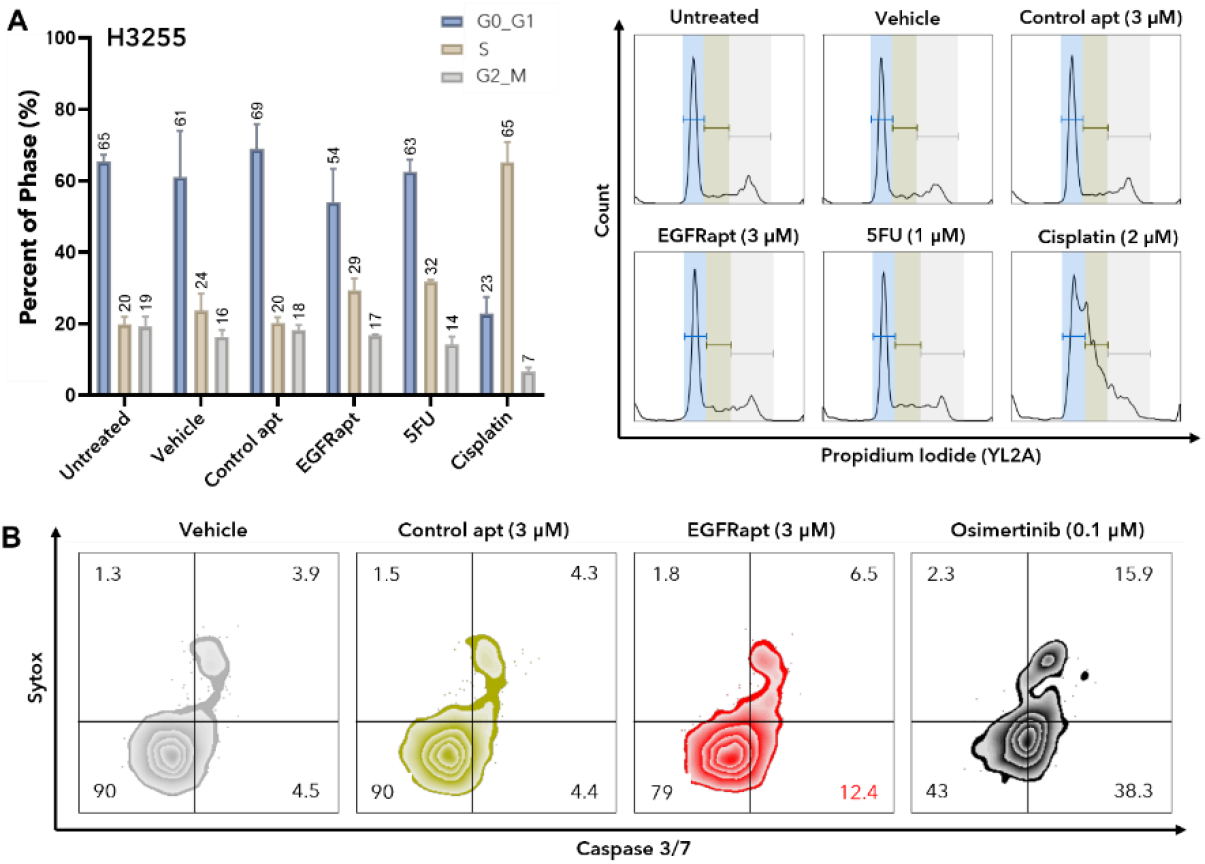
*EGFRapt* induces cell cycle arrest and apoptosis in H3255. (**A**) H3255 cells were treated with reported concentrations of *EGFRapt*, control apt, or chemotherapeutic for 48 h, then fixed and stained with propidium iodide. Cell cycle determination was analyzed by flow cytometry and using the Watson pragmatic method. The percentage of cells in the G0-G1 (blue bar), S (sand bar), or G2-M (grey bar) phase of cell cycle is plotted on the *y*-axis. Plotted values represent mean ± SD for n=2-3 independent experiments. H3255 cells treated with *EGFRapt* increased percentage of cells in the S phase compared to non-binding control apt or vehicle, similar to S phase specific inhibitor 5FU. Representative histograms with fill colors corresponding to phase of cell cycle are shown on the right. (**B**) H3255 cells were treated as in A but were live stained with apoptosis marker (CellEvent^TM^ Caspase 3/7 reagent; *x*-axis) and cell impermeable necrosis and late apoptosis marker (Sytox^TM^, *y*-axis). Cell staining was analyzed by flow cytometry. Representative contour plots are shown in which percent of cells unstained (Q3) or stained with Caspase 3/7 reagent only (Q4), Sytox^TM^ only (Q2), or both reagents (Q1) from one of n=2 independent experiments. H3255 cells treated with *EGFRapt* increased percentage of cells stained with Caspase 3/7 reagent when compared to control apt or vehicle.

### EGFRapt is anti-proliferative in H1975 but not H3255

Given that H1975 cells exhibited only a partial, non-cytotoxic response to *EGFRapt* and that brightfield images of the dose response hinted at different phenotypes between H1975 and H3255 (Figure 1E), we speculated that either the mechanism of *EGFRapt* in H1975 differs from that of H3255 or that H1975 harbors a mechanism of targetable resistance to aptamer treatment. To determine whether EGFR kinase-dependent functions were altered by treatment with *EGFRapt*, we monitored proliferation via carboxy fluorescein succinimidyl ester (CFSE, tracking dye for viable cells) and Ki67 (a cellular protein marker of proliferation) staining, and canonical signaling pathways via western blot. CFSE and Ki67 staining revealed a decrease in proliferation upon treatment with a single dose of 3 µM *EGFRapt* when compared to vehicle, or when compared to a non-binding or binding control apt (**Figure 4A-B**). The western blot of phosphorylated and total EGFR and downstream signaling partners, ERK and AKT, revealed that a single dose of *EGFRapt* decreased phosphorylation levels of ERK1/2 but intriguingly had little to no effect on phosphorylation of EGFR or AKT. The magnitude was further amplified upon a second dose of *EGFRapt* (**Figure 4C**). Notably, the effect of *EGFRapt* on H1975 signaling was different from that of other EGFR therapeutics, Osimertinib (TKI) and Nimotuzumab (mAb). Consistent with previous reports^52,53^, Osimertinib treatment decreased phosphorylation of EGFR, ERK1/2, and AKT whereas Nimotuzumab decreased phosphorylation of EGFR and ERK1/2 and decreased total EGFR (**Figure 4C**), likely due to induced receptor internalization and degradation. Treating H3255 cells with a single dose of 3 µM *EGFRapt* did not lead to similar differences in proliferation (CFSE, **Figure S6A**) or canonical signaling pathways (**Figure S6B**); however, two doses of 3 µM aptamer did decrease phosphorylation of EGFR but not downstream signaling partners (ERK1/2 or AKT). Taken together, these findings suggest that *EGFRapt* is anti-proliferative in H1975 and are consistent with alteration of the canonical, kinase dependent functions of EGFR, a mechanism different from H3255.

**Figure 4.**
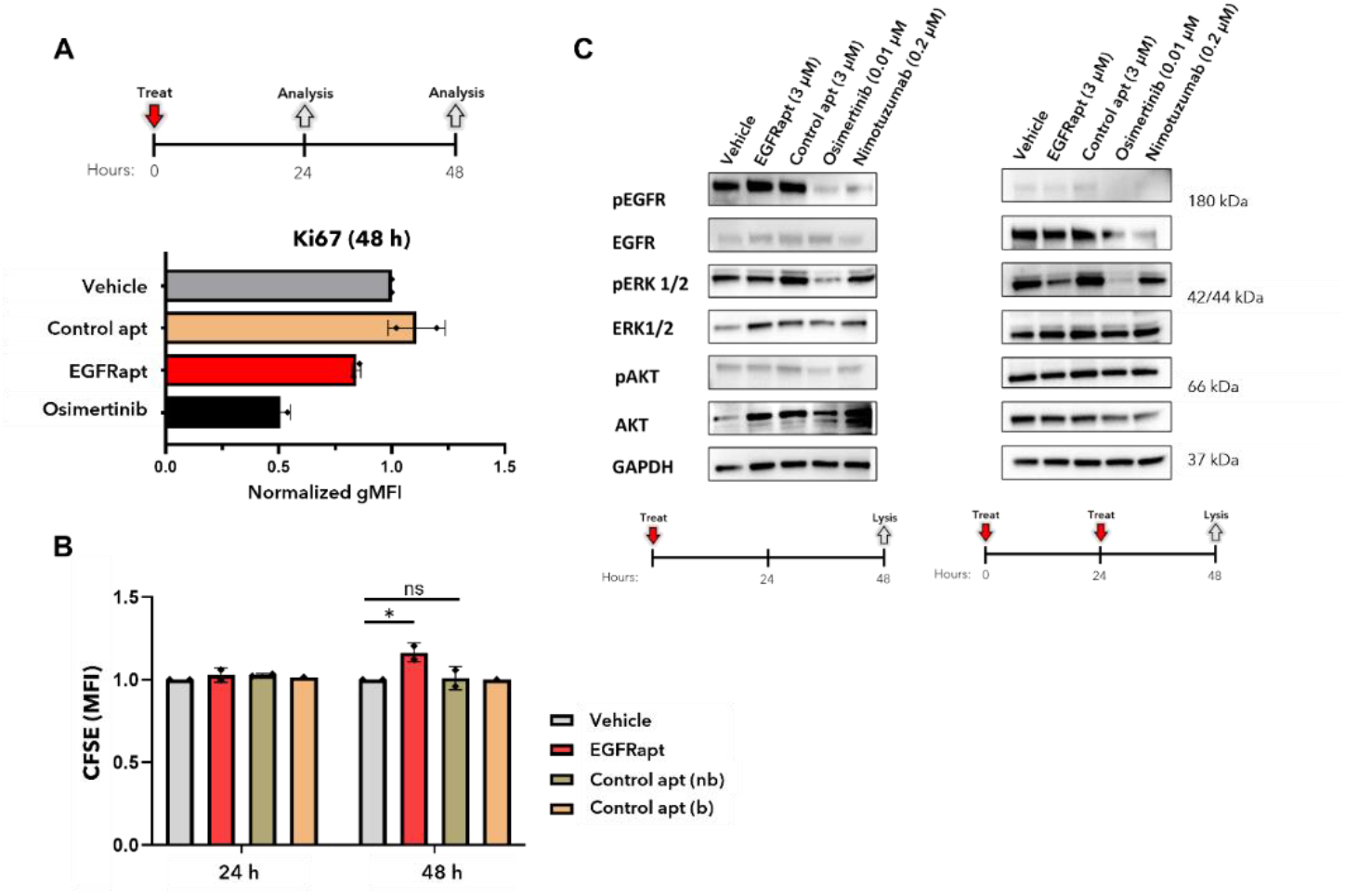
*EGFRapt* alters canonical EGFR signaling and is anti-proliferative in H1975. (**A-B**) H1975 cells were prelabeled with Carboxy Fluorescein Succinimidyl Ester (CFSE) and treated with 3 µM *EGFRapt* (red bars), binding (b, orange bars) or non-binding (nb, tan bars) control apts, 0.1 µM Osimertinib (black bar), or vehicle (grey bars) for 24 or 48 h and then fixed and stained with APC labeled Ki67 antibody. CFSE and antibody staining were analyzed by flow cytometry. The mean fluorescence intensities (MFI) of Ki67 and CFSE staining are plotted on the *x-*axis and *y-*axis, respectively. H1975 cells treated with *EGFRapt* exhibited decreased Ki67 staining and increased CFSE staining at 48 h, both suggestive of decreased proliferation. (**C**) Total cell lysates from H1975 cells treated with reported doses of one (*left*) or two (*right*) doses of *EGFRapt*, non-binding control apt or vehicle for 48 h, one dose of Nimotuzumab for 48 h, or one dose of Osimertinib for 16 h were probed for total and phosphorylated EGFR, ERK1/2, AKT, or loading control, GAPDH. Quantification of blot intensity (data not shown) revealed decreased phosphorylation of ERK1/2 in *EGFRapt* but not control apt or vehicle treated cells.

### Bulk RNA sequencing analysis supports anti-cancer mechanisms in H3255 and H1975

As an independent assessment of the anti-cancer mechanism of action for *EGFRapt*, we preformed post hoc bulk RNA sequencing for all four NSCLC cell lines (H3255, H1975, H820, and A549) treated with 3 µM *EGFRapt*, control apt, or vehicle for 24 h. Principal component analysis (**Figure S7A**) and FPKM normalization with Pearson correlation-based clustering of gene expression (similarity matrix, **Figure S7B**) revealed tight, cell line-based clustering with minimal overlap among the four cell lines regardless of treatment, suggesting minimal overall transcriptional reprogramming. Based on differential expression analysis, H3255 and H1975 cells treated with *EGFRapt* relative to control apt had 1002 (801 unique) and 520 (304 unique) differentially expressed genes (DEGs; |Log2FC| ≥ 0.5 and *q*-value ≤ 0.05), respectively (**Figures 5A and S7C-D**), with only 90 DEGs being shared between the two cell lines (**Figure 5A-B**). In contrast, cell lines A549 and H820, which had shown no effect in cell viability upon *EGFRapt* treatment (Figure 1A), had much fewer DEGs (196 and 206, respectively) when comparing *EGFRapt* to control apt (**Figures 5A and S7C-D**). These high-level RNA sequencing data highlight the minimal overlap in the changes of gene expression among cell lines upon treatment with *EGFRapt* and is consistent with the anti-cancer mechanisms being cell specific.

**Figure 5.**
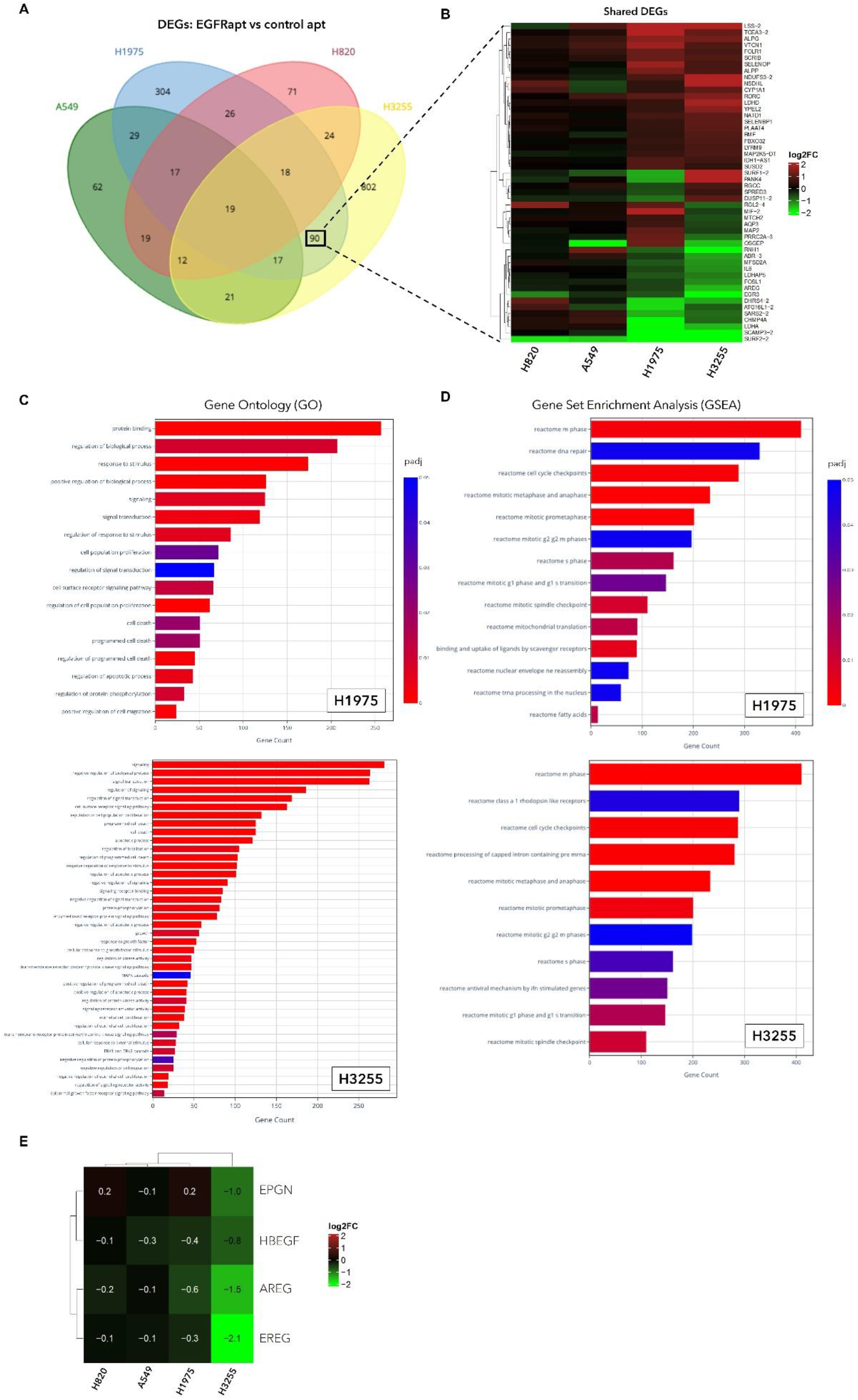
Pathway analyses of differentially expressed genes in *EGFRapt* treated cells independently supports mechanistic studies. (**A-E**) H3255 and H1975 cells were treated with 3 µM *EGFRapt* or control apt for 24 h. Cells were then lysed and total RNA was subjected to bulk RNA sequencing and analysis. (**A**) Venn-diagram of number of differentially expressed genes (DEGs) for all cell lines treated with EGFRapt compared to control apt. (**B**) Heatmap showing relative expression of DEGs shared by H3255 and H1975. (**C**) Gene ontology terms associated with biological process (GO:BP) and molecular function (GO:MF) in H1975 (*top*) and H3255 (*bottom*) treated with *EGFRapt* compared to control apt. (**D**) Gene set enrichment analysis (GSEA) terms associated with Reactome pathways in H1975 (*top*) and H3255 (*bottom*) treated with *EGFRapt* compared to control apt. (**E**) Heatmap showing relative expression of four EGFR ligands (epithelial mitogen, heparin-binding EGF-like growth factor, amphiregulin, epiregulin). For B and E, red = upregulated; green = downregulated; black = no change. *EGFRapt* induced notably more gene expression changes in H3255 and H1975 compared to A549 and H820; these changes were mostly cell line dependent with only 90 DEGs being shared between H3255 and H1975. Pathway analyses (GO and GSEA) in both cells lines treated with *EGFRapt* compared to control apt revealed significant enrichment of terms relating to processes identified through experimental findings from Figures 3 and 4, such as receptor signaling, proliferation, programmed cell death, and cell cycle phase transition.

Pathway enrichment analyses from cells treated with *EGFRapt* relative to control apt are shown in **Figure 5C-D**. Significantly enriched (*q*-value ≤ 0.05) Gene Ontology (GO) terms associated with Biological Processes (BP) or Cellular Component (CC) were identified using DEGs with GProfiler. DEGs associated with each GO term for all cell lines (H3255, H1975, A549, H820) are shown in the *supplementary data tables*. Significantly enriched Reactome and KEGG terms were identified using weighted and ranked expression changes (regardless of log2FC) with Gene Set Enrichment analysis (GSEA). For both H3255 and H1975, significant GO:BP terms were linked to receptor signaling, signaling transduction, proliferation, and programmed cell death (**Figure 5C**), and significant Reactome and KEGG terms were linked to the cell cycle and its checkpoints (**Figure 5D**). Nearly all enriched Reactome terms harbored negative normalized enrichment scores, suggestive of an overall downregulation of genes involved in these pathways (**Figure S8A**). We also observed that for both cell lines, enriched GO:CC terms suggested similar subcellular localization (**Figure S9A-B**) with enriched terms including ‘membrane’, ‘extracellular vesicle’, ‘membrane bound organelle,’ and ‘endosome’. Comparing H3255 and H1975 at this level suggests that many similarities are present between the two cell lines upon *EGFRapt* treatment. These results are not unexpected, as canonical signaling partners of EGFR (e.g., ERK1/2)^54^ and non-canonical functions of EGFR (e.g., nuclear translocation)^12,13^ have been shown to regulate both cell cycle progression and proliferation. Looking in more depth at these data and the relative expression of genes involved in cell cycle related Reactome terms, H3255 exhibited a notably larger change in gene expression compared to H1975 (**Figure S8B**). This could explain why H3255 cells were more sensitive in cell cycle analysis (Figure 3A). In addition to the number of significant GO:BP pathways being much larger in H3255 compared to H1975, the number of DEGs involved in many of these pathways was also notably larger (Figure 5C and *supplementary tables*). This includes pathways related to programmed cell death and apoptosis, processes that were shown to be involved in the anti-cancer mechanism of H3255 (Figure 3B). Interestingly, multiple pathways regarding regulation of signaling were significantly enriched, including some relating to canonical EGFR signaling (terms: ‘EGFR signaling pathway’, ‘MAPK cascade’, ‘ERK1/2 cascade’), despite minor changes seen at the protein level (Figure S6B). Within these pathways, the expression of four EGFR ligands – *EPGN, HBEGF, EREG, and AREG* – were remarkably downregulated in only H3255 cells (**Figure 5E**). As these ligands are involved in both EGFR homodimer and heterodimer signaling, these gene expression changes may also play roles in the differential cell fates upon treatment with *EGFRapt*.

Overall, bulk RNA sequencing analysis generally supports the anti-cancer mechanisms observed above (i.e., anti-proliferative in H1975 or cytotoxic in H3255); however, this analysis also supports that while the interaction between *EGFRapt* and EGFR may be similar between the two cell lines, their fates are highly dependent on the gene expression changes (Figures 5, S7 and S8) and protein interactions (Figures 4C and S6B) that occur downstream. As H3255 had shown a complete response and H1975 a partial response to treatment with *EGFRapt,* better understanding of how these gene expression changes impact cell survival will be important to understanding the sensitivity to *EGFRapt,* its highly modified variants, and potentially other EGFR inhibitors.

### EGFRapt decreases cell surface expression of EGFR but the anti-cancer function of EGFRapt is not affected by inhibitors of endocytosis

Anti-EGFR mAb treatment has in part shown efficacy in EGFR positive tumors due to their ability to internalize the receptor and mark it for degradedation^52,55^. As bulk RNA sequencing analysis identified enrichment of GO:CC terms relating to the cell membrane and vesicular compartments, we sought to determine whether surface expression of EGFR was similarly altered after treatment with *EGFRapt*. Treatment with *EGFRapt* decreased cell surface expression of EGFR at 24 h (and to a lesser extent at 48 h) in both H1975 and H3255, as determined by MFI of a non-competing, APC-labeled anti-EGFR antibody (clone AY13)^47^ was probed at 4°C to minimize recognition of internalized receptor (**Figure S10A**). However, unlike anti-EGFR mAbs, immunoblotting of total protein on SDS-PAGE gels showed similar levels of total EGFR before and after treatment with *EGFRapt* (**Figure 4C** and **S10A**), indicating that the observed decrease in cell surface expression is not associated with degradation of the receptor, but is instead potentially due to increased internalization or decreased recycling of the receptor back to the cell surface after treatment.

As cetuximab has been reported to internalize EGFR in a tyrosine kinase-independent manner and at a much slower rate than ligand EGF^56^, we next sought to determine whether clathrin mediated endocytosis (CME; a major mechanism of tyrosine kinase-dependent and ligand stimulated EGFR internalization) or clathrin-independent endocytosis (CIE; a minor mechanism of EGFR internalization) played a significant role in the mechanism of *EGFRapt*. Therefore, we assessed cell viability of H3255 cells treated with 3 µM *EGFRapt* in the presence or absence of three separate endocytosis inhibitors—methyl β cyclodextrin (MβCD; cholesterol depletion in lipid rafts affecting mostly CIE but also CME), Pitstop 2 (clathrin inhibition affecting mostly CME but also CIE), and Dynasore (dynamin inhibition affecting dynamin-dependent CME and CIE). None of the endocytosis inhibitors significantly diminished the anti-cancer effect of *EGFRapt* relative to the effects observed when *EGFRapt* was administered alone, even at inhibitor concentrations that negatively impacted cell viability in the absence of *EGFRapt* (**Figure S10B-C**). These data suggest that the mechanism by which *EGFRapt* acts is likely independent of clathrin, dynamin, and cholesterol mediated endocytosis; however, as endocytosis inhibitors have consistently shown lack of specificity^57,58^, we cannot completely rule out dependence on these forms of endocytosis.

### EGFRapt combined with chemotherapeutics or tyrosine kinase inhibitor increases anti-cancer effect in H1975

NSCLC treatment regimens are typically adjusted based on disease progression or mutation profiles obtained from sequencing of the tumor specimen or circulating tumor DNA. These changes include the substitution or addition of targeting agents and salvage chemotherapy. Both Osimertinib and *EGFRapt* were shown above to prevent phosphorylation of ERK1/2, albeit via different upstream mechanisms, raising the possibility that both compounds could be used together. To determine whether *EGFRapt* was still efficacious in or provided additional benefit to chemotherapeutics or TKIs, we treated H1975 cells with *EGFRapt* in combination with these therapies. Treatment of 3 µM *EGFRapt* with near or sub-EC50 concentrations of doxorubicin (5 µM), cisplatin (10 µM), or Osimertinib (0.001 µM) further decreased cell viability compared to these therapies alone (**Figure 6A**). A dose response with Osimertinib revealed that the addition of *EGFRapt* only decreased cell viability when using sub-IC50 concentrations of Osimertinib (**Figure 6B**). In general, these data suggest that *EGFRapt* could provide additional benefit when used in conjunction with chemotherapeutics or TKIs (*e.g.*, adjuvant therapy).

**Figure 6.**
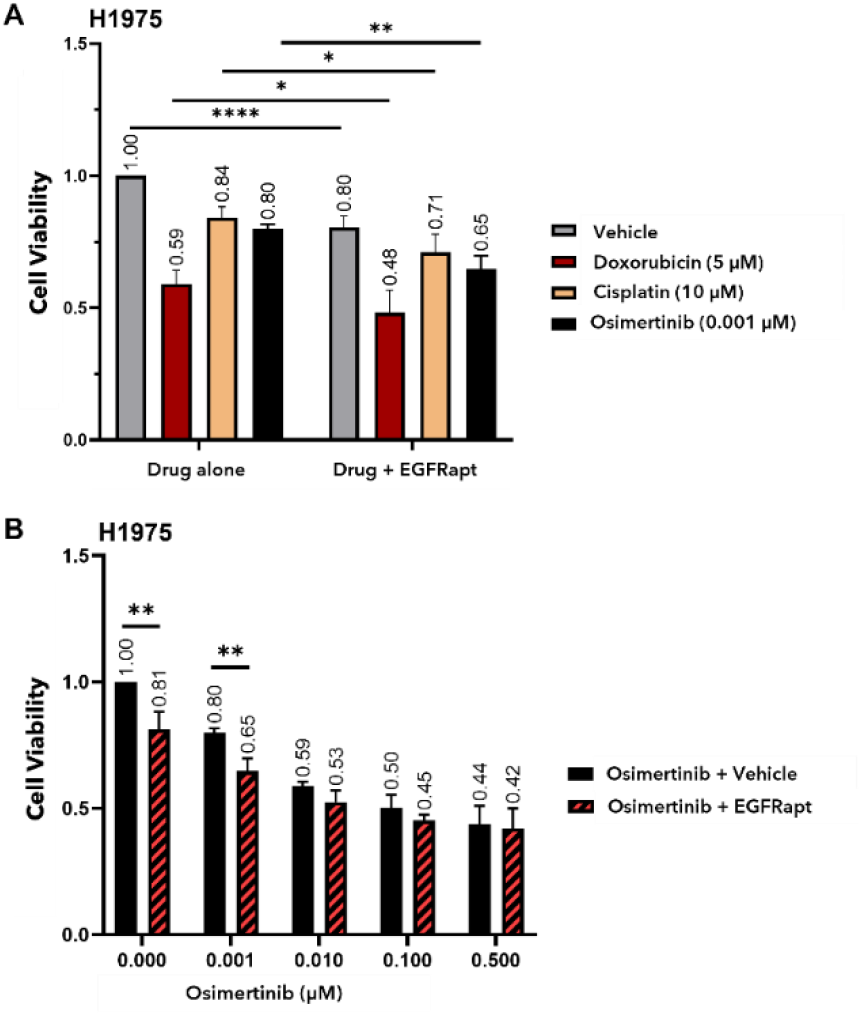
Combining *EGFRapt* with chemotherapeutics or TKI enhances anti-cancer effect in H1975. (**A-B**) H1975 cells were subject to cell viability assay as in Figure 1B except that cells were treated with a combination of 3 µM *EGFRapt* or vehicle and reported concentrations of three clinically relevant anti-cancer reagents—Doxorubicin (dark red bar; EC50 ∼5 µM), Cisplatin (orange bar; EC50 ∼20 µM), or Osimertinib (black bar; EC50 ∼0.5 µM)—or vehicle (grey bar). EC50 values for this cell line were determined using cell viability of H1975 at 48 h in response to drug only (data not shown). Cell viability was determined using MTS reagent. Combination treatment decreased cell viability compared to anti-cancer reagent alone (A). For Osimertinib, combination treatment (red/black bar) decreased cell viability compared to Osimertinib alone (black bar) only at sub-EC50 concentrations (B). For A-B, relative cell viability (normalized to vehicle) is reported on the y-axis and plotted values represent mean ± SD for n=2-3 independent experiments, each with 2-4 technical replicates. Statistical analysis was preformed using a two tailed t-test (*p < 0.05, **p < 0.01, ****p < 0.0001).

## Discussion

Poor survival rates reported under the current therapeutic landscape for patients with NSCLC harboring mutant EGFR warrant the development of novel therapeutics with unique mechanisms of action. Here we showed that *EGFRapt* and heavily modified versions of *EGFRapt* (Figure S4) provide a novel means of targeting and treating NSCLC cell lines and tumors that harbor mutant EGFR, especially when delivered optimally over multiple doses or in combination with chemotherapeutics or TKIs. Furthermore, preliminary evaluation of the anti-cancer mechanisms by which *EGFRapt* acts in two different cell lines harboring L858R ± T790M mutations indicate cell cycle arrest and apoptosis in cell line H3255 and preventing phosphorylation of downstream signaling partners to slow proliferation in cell line H1975 (**Figure 7A**). Importantly, these mechanisms differ from that of current FDA approved treatments. In NSCLC, oncogenic EGFR TKD mutations typically drive uncontrolled canonical signaling through ‘inside-out’ activation (*i.e.*, dimerization and autophosphorylation without ligand stimulation)^59,60^. TKIs usually inhibit these canonical functions by competing for ATP binding sites in the intracellular TKD to prevent dysregulated kinase domain phosphorylation and downstream signaling cascades^61^. Conversely, anti-EGFR mAbs (*e.g.*, cetuximab, panitumumab, necitumumab, and nimotuzumab) bind to the extracellular ligand binding or dimerization domains of EGFR and block ligand interaction or prevent EGFR dimerization^62–65^, and/or induce receptor internalization and degradation^52,55^. As these interactions prevent ‘outside-in’ activation, mAbs are often, but not always^66–69^, ineffective at treating NSCLC harboring mutant EGFR^70,71^, for which signaling occurs independent of ligand stimulation. Nonetheless, the mechanisms of both TKIs and mAbs are dependent directly on decreasing the number of phosphorylated and actively signaling EGFRs. Here we show that, for these cell lines that express the L858R activating mutation, *EGFRapt* does not similarly alter pEGFR (or total EGFR) levels but does affect surface expression of EGFR and downstream canonical signaling partners in H1975 or non-canonical functions in H3255.

**Figure 7.**
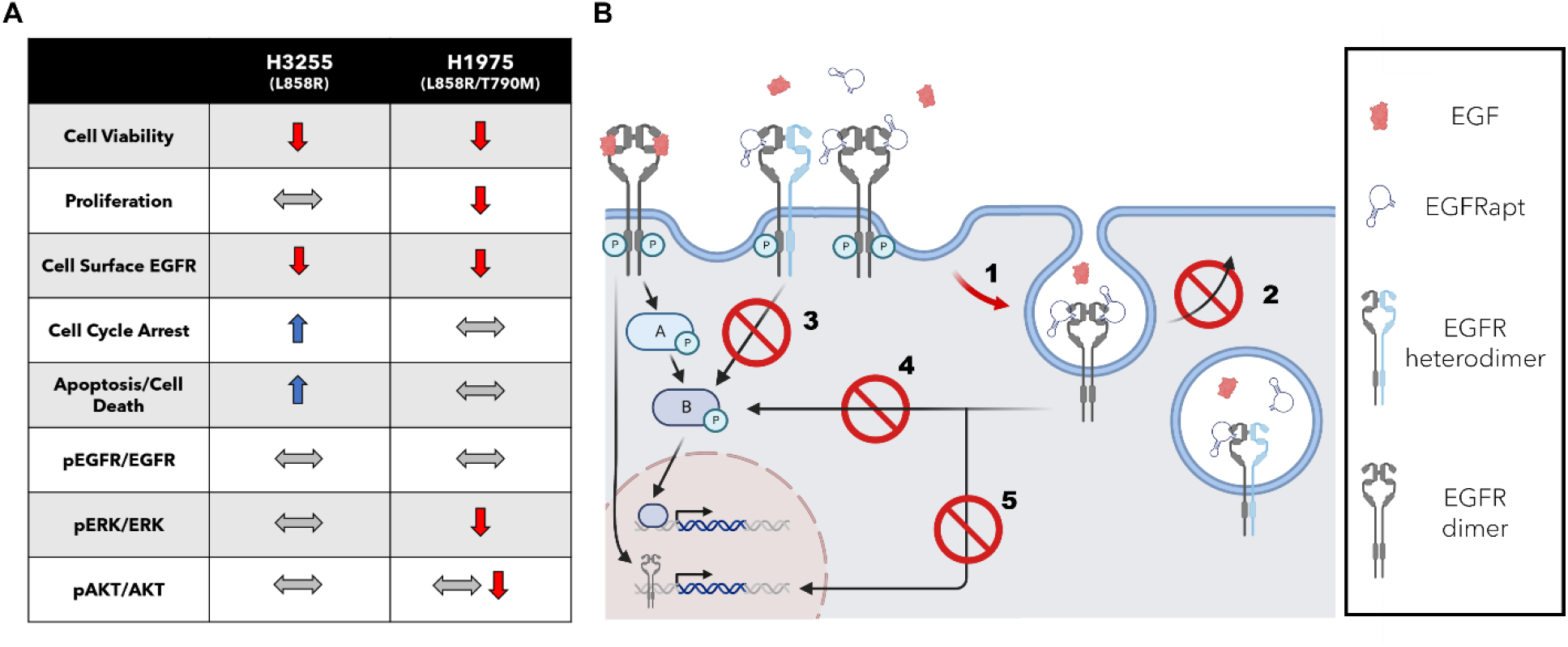
Summary of *EGFRapt* effects on NSCLC cell lines harboring L858R mutations. (**A**) Two cell lines harboring EGFR L858R plus (H1975) or minus (H3255) T790M mutation exhibited decreased cell viability upon treatment with *EGFRapt* when compared to control apts or vehicle. H1975 cells treated with *EGFRapt* exhibited impaired proliferation consistent with alteration of kinase-dependent (canonical) EGFR signaling. H3255 cells treated with *EGFRapt* exhibited increased cell cytotoxicity and disruption of cell cycle phase transition consistent with alteration of kinase-independent (non-canonical) EGFR signaling. (**B**) Hypothesized anti-cancer mechanisms of *EGFRapt* in H1975 and H3255. EGFR internalization (1) or recycling (2) is altered upon *EGFRapt* binding, leading to lower cell surface expression. In H1975, EGFR mistrafficking, along with possible disruptions of EGFR heterodimer formation, prevents activation of downstream signaling partners (3 and 4), leading to decreased proliferation. In H3255, EGFR mistrafficking prevents non-canonical functions that promote survival, such as those driven by nuclear localization of EGFR (5).

*EGFRapt* has previously been shown to have anti-proliferative properties in an epidermoid cancer cell line, A431^43^, which expresses high levels of WT receptor. However, as *EGFRapt* was reported to compete with EGF^43^ and binds without disrupting EGFR homodimerization^47^ (*i.e.*, it does not disrupt the dimerization domain), it is likely the anti-cancer mechanism in A431 is unrelated to the mechanisms we describe here for cell lines harboring mutant EGFR (H3255 and H1975) but is instead similar to that of mAbs, in that they prevent ligand-induced, ‘outside-in’ activation. The lack of significant inhibition of mutant EGFR phosphorylation via this mechanism in our study is consistent with ‘inside-out’ receptor activation (due to the L858R mutation causing constitutive activation in the absence of a stimulating ligand). Still, we cannot rule out, especially for multiple doses of *EGFRapt in vivo*, whether additional mechanisms may contribute to anti-cancer effects, such as competing with and preventing ligand stimulation.

Drug resistance is a major contributing factor to treatment failure and disease progression, and sub-optimal concentrations are known to provide selective and/or adaptive pressures for tumor evolution. Dosage strategies (*i.e.*, amount and timing) have been shown to be effective in preventing or delaying acquired resistance^72^. Combination therapy is an alternative method to overcome the drug resistance that is typically seen in sequential monotherapy treatment^72–75^ and has the added benefit of enhanced tumor killing. In the era of personalized medicine, clinical trials using a combination of therapeutic interventions are increasing. In this work we showed that *EGFRapt* could be used in combination with other therapeutics to further decrease cell viability in H1975 cells (Figure 5). In addition to resistance, dose limiting adverse events (AEs) are another major contributing factor to treatment failure and have been shown to severely affect patients’ quality of life. The most commonly reported TKI induced AEs are grade 1-3 cutaneous and mucosal reactions^76,77^, although more severe and even fatal AEs have been reported.^77^ Use of *EGFRapt* in the adjuvant or neoadjuvant setting with TKIs could be advantageous, as it may allow lower doses of each drug be used to elicit a similar anti-cancer effect, and this may minimize associated adverse events (AEs) and improve quality of life.

Most of the studies reported in this work utilized a partially modified (2’FY) and unconjugated version of *EGFRapt*. Unfortunately, such aptamers are metabolically unstable and are susceptible to significant nuclease degradation and excretion from circulation within a few hours of administration^37,78^. The two currently FDA approved therapeutic aptamers are extensively modified post-selection (including PEGylation) and are delivered locally (*i.e*., intraocular injection), rather than systemically^79,80^. The present work includes an initial exploration of highly modified versions of *EGFRapt* in which ∼85% of the nucleotides carry 2’FY or 2’OMeR modifications (Figure S4). The original aptamer selection was carried out using 2’FY and fully 2’OH purines, and the aptamer appears to require that some of the purine positions retain the hydroxyl^51^. Nevertheless, applying additional medicinal chemistry and molecular engineering methods to address pharmacokinetic barriers will be important to the reagent’s success *in vivo* and ultimately for clinical translation. Alternatively, improving methods for local delivery to tumors could provide strategies to increase concentrations of the reagent at the tumor. Methods such as intranasal drip or inhalation have been successfully applied for delivery of oligonucleotide therapeutics, including aptamers, siRNAs, and ASOs, to the lung^81,82^. In cases where partially modified and unconjugated oligonucleotides with less favorable pharmacokinetics and biodistribution profiles must be used, local delivery may expose the reagent to fewer serum nucleases and provide high local tissue distribution before passing through the kidney filtration system. Our subcutaneous human xenograft model, where tumors were soft, palpable, and close to the surface, made localizing the tumor and injecting our aptamer intratumorally relatively easy, in contrast to some fibrotic (*i.e.,* impenetrable) or deep tissue tumors. In a more clinically relevant, orthotopic model, intratumoral delivery becomes much more challenging, and metastasis outside of the lung is common in late-stage disease; therefore, a mix of local delivery and systemic administration of highly modified aptamers may be best.

Our bulk RNA sequencing analysis not only supports experimental findings but also positions us to speculate on the upstream mechanisms that led to the anti-cancer processes described above. Regulation of EGFR signaling through heterodimerization or crosstalk with other proteins, non-receptor- and receptor tyrosine kinases (RTKs) has been readily reported, especially with other ErbB-family members (*e.g.*, HER2)^83–88^. Aberrant trafficking of RTKs has also been shown to significantly impact their signaling, especially when they colocalize with other receptors^16,83,89,90^. As significantly enriched GO:BP terms in both H3255 and H1975 suggested modulation of receptor signaling and signaling transduction, it is plausible that treatment with *EGFRapt* disrupts EGFR heterodimerization and/or receptor crosstalk with other proteins, either on the cell surface or in intracellular compartments such as endosomes. Cyclooxgenase-2 (COX-2/PTGS2) is one such protein that has been shown to participate in crosstalk with EGFR to orchestrate tumor growth^91^. Multiple mechanisms have been cited, including modulating the production of proteins and lipids (e.g., PGE_2_) that regulate the expression of EGF-like ligands (e.g., amphiregulin, as we have seen in H3255; Figure 5E) or alter phosphorylation of proteins that act downstream of EGFR (e.g., ERK, as we have seen in H1975; Figure 4C). Interestingly, our bulk RNA sequencing analysis revealed that *PTGS2* expression was significantly downregulated in both H1975 and H3255 upon treatment with *EGFRapt* (Figure S11A). As multiple reports have shown that cell surface^91^ and nuclear^13^ EGFR regulates COX-2 expression through interactions with STAT5 and the transcription factor STAT3, respectively, it is tempting to speculate that *EGFRapt* treatment is decreasing *PTGS2* expression in an EGFR dependent manner, which is in turn further negatively impacting EGFR function (i.e., cyclic regulation). However, such speculation will require further exploration and validation prior to defining involvement of this pathways in the response to *EGFRapt* treatment.

Our findings in H3255 were highly suggestive that pro-survival (non-canonical) functions are disrupted upon *EGFRapt* treatment, as such treatments did not alter phosphorylation of downstream signaling partners but did disrupt cell cycle progression and induce apoptosis. In addition to cell cycle disruption, we also noted a marked down regulation of genes involved in cytokine/chemokine and type I interferon signaling (*CCL2/22, IFIT1/2/3, ISG15, CXCL1/2/8,* and *IL6*) in H3255 treated with *EGFRapt* versus control apt or vehicle (**Figure S11A-B**). Importantly, the type I interferon response has been reported to be both pro-survival (tumorigenic) and cytotoxic (anti-tumor), depending on the cancer type^92^. Induction of a type I IFN response upon treatment of mutant EGFR NSCLC with TKIs has been shown to cause resistance, and inhibition of IFN beta (a type I IFN) was shown to restore TKI efficacy and anti-tumor responses^93,94^. Furthermore, cancer cells with p53 mutations, such as those found in H3255^95^, have also been shown to rely on type I IFNs for survival. As L858R mutations are known to impair EGFR nuclear transport^96^, it is plausible that treatment with *EGFRapt* further disrupts intracellular trafficking of EGFR to the nucleus and impairs its nuclear functions in gene expression and DNA repair^12^. This is consistent with our findings of decreased surface expression, impaired cell cycle progression, and enrichment of GO or GSEA terms relating to regulation of localization and cell cycle disruption (Figure 5C-D), but this model is speculative and requires further confirmation. Nevertheless, taken together our findings from transcriptomics analysis in H3255 and H1975 provide insight into additional mechanisms that could potentially be targeted in NSCLC harboring L858R ± T790M mutations and that warrant further investigation.

Overall, our findings show that *EGFRapt* is a promising reagent for continued therapeutic development and could potentially provide an alternative strategy to treat patients with mutant EGFR positive NSCLC. As primary and secondary mutations in EGFR have been shown to rewire the receptors interactome and alter intracellular trafficking^14,17,97^, efforts are ongoing to further define the mechanism of action of *EGFRapt* in these NSCLC cell lines (**Figure 7B**) that may be targetable by other means (e.g., COX-2 inhibition using NSAIDs), to elucidate whether the mechanisms are L858R mutation specific, and to establish whether consequences of the secondary T790M mutation, or an alternative mechanism, is responsible for the incomplete anti-cancer response seen in H1975. It will be especially revealing to determine whether patient tumor samples respond similarly. Furthermore, as anti-EGFR mAbs have efficacy in other cancer types such as colorectal and head-and-neck cancer, and *EGFRapt* binds epitopes that overlap with these mAbs, efforts are also ongoing to determine whether *EGFRapt* has anti-cancer activity in these cancer types.

## Supporting information

RNAseq data tables

## Acknowledgements

This work was financially supported by the MU Life Sciences Center (LSC) - Early Concept Grant (ECG) for Innovative Collaborative Research involving Post-Doctoral Researchers (PI: Porciani-Daniels-Burke), UM Research and Creative Works Strategic Investment Program Grant (PI: Burke) and Missouri Department of Health and Senior Services (MDHSS) - Contract #AOC23380006 (PI: Joshi).

## CRediT authorship contribution statement

**Brian J. Thomas:** Conceptualization, Methodology, Investigation, Visualization, Writing – Original Draft, Writing – Review & Editing. **Sania Z. Awan:** Methodology, Investigation, Visualization, Writing – Review & Editing. **Trupti Joshi:** Resources, Supervision, Writing – Review & Editing. **Mark A. Daniels:** Resources, Supervision, Funding acquisition. **David Porciani**: Conceptualization, Visualization, Writing – Review & Editing, Supervision, Funding acquisition. **Donald H. Burke:** Project administration, Writing – Review & Editing, Supervision, Funding acquisition.

## Data and code availability statement

The dataset used in the RNA sequencing analysis has been deposited in NCBI Gene Expression Omnibus (GEO) Database, where it is accessible under accession number GSE259407. For interactive analysis, the analyzed dataset can be found at KBCommons (https://kbcommons.org/system/browse/diff_exp/homoSapiens), a cutting-edge framework and visualization tool, which integrates an extensive suite of bioinformatics tools.

## Materials and Methods

### Reagents

DNA oligonucleotides were purchased from Integrated DNA Technologies (IDT, Coralville, IA). Modified aptamers that carried both 2’FY and 2’OMeR were purchased from TriLink BioTechnologies (San Diego, CA). APC-labeled EGFR monoclonal antibody (clone AY13, cat: #352906), Ki67 (cat: # 350514), and isotype control (mouse IgG1, κ, clone MOPC-21, cat: #400120) were purchased from BioLegend (San Diego, CA). EGFR monoclonal antibodies (cetuximab and nimotuzumab) were purchased from Novus Biotechnology (Centennial, CO). Primary antibodies for probing western blots were purchased from Cell Signaling Technology (Danvers, MA; (p)EGFR), ABclonal Technology (Woburn, MA; (p)AKT, (p)ERK1/2), and BioLegend (GAPDH, cat #607902). Secondary antibodies for western blots were purchased from Invitrogen (Waltham, MA, cat #G-21040, #G-21234, #A18871). Dual color protein ladder standards were purchased from Biorad (cat #1610374). BSA standards and Pierce detection reagents were purchased from ThermoFisher Scientific (Waltham, MA). Protease and phosphatase inhibitor cocktails and RIPA buffer were purchased from Abcam (Cambridge, MA). All other reagents and materials, including chemotherapeutics and small molecule inhibitors, were purchased from Sigma Aldrich (St. Louis, MO) unless otherwise noted.

### Aptamer generation

All 2’FY-modified RNA and DNA aptamer sequences are reported in Table S1, with additional detail in Figure S4. All DNA template and primer oligonucleotides (Table S2) were resuspended in appropriate volume of Milli-Q® water to reach a stock concentration of 100 µM. 2’FY modified RNA aptamers were generated via *in vitro* run-off transcription (IVT). First the DNA templates were PCR amplified with primers that appended a T7 promotor (Table S3). Then, 2′FY-modified RNA aptamers were transcribed by overnight IVT at 37°C using recombinant mutant T7 RNA polymerase (Y639F), IVT buffer (50 mM Tris-HCl pH 7.5, 15 mM MgCl_2_, 5 mM DTT, 4% w/v PEG4000 and 2 mM spermidine), and 2 mM each of ATP, GTP, 2’-fluoro modified CTP and 2’-fluoro modified UTP (TriLink Biotechnologies). All RNA aptamers were purified through denaturing polyacrylamide gel electrophoresis (0.75 mm, 6-8% TBE-PAGE, 8 M urea) and bands corresponding to the expected product sizes were visualized by UV shadow, excised from the gel, and then eluted overnight while tumbling in 300 mM sodium acetate pH 5.4. Eluates were ethanol precipitated, resuspended in Milli-Q® water, and stored at −20°C until further use. If concentrating was required, 2’FY RNA aptamers were lyophilized and resuspended in an appropriate volume of Milli-Q® water.

### Aptamer folding

All aptamers were folded as previously described^29,48^. In brief, *in vitro* folding reactions were prepared at room temperature in Dulbecco’s phosphate buffered saline (DPBS, pH 7.4) supplemented with 5 mM MgCl_2_. For thermal renaturation, samples were transferred into a preheated aluminum insert within a dry heat block set to 90°C, where they were kept for 2-3 min to denature nucleic acid structures, and then the aluminum insert was removed from the block heater and placed on the workbench to cool slowly to below 37°C. For *in vitro* applications, aptamer samples were freshly prepared before each experiment at 10X concentration and diluted to 1X in complete media. For *in vivo* intratumoral injections, aptamer was prepared at 3 µM in DPBS without MgCl_2_.

### Cell culture

A549 cells were gifted by Dr. Bumsuk Hahm, University of Missouri-Columbia. H3255, H1975, and H820 were gifted by Dr. Raghuraman Kannan, University of Missouri-Columbia. MCF7 cells were gifted by Dr. Thomas Quinn, University of Missouri-Columbia. A549 and MCF7 cells were cultured in DMEM supplemented with 10% Fetal Bovine Serum (FBS), 1 mM sodium pyruvate, 2 mM L-glutamine, and 1X NEAAs. H1975, H3255, and H820 cells were cultured in HyClone™ RPMI 1640 supplemented with 10% FBS, 1 mM sodium pyruvate, 2 mM L-glutamine, and 1X NEAAs. All cell lines were maintained at 37°C in humidified incubator with 5% CO_2_ and passaged when ∼80% confluent. Cells were not passaged more than 20 times from thaw or past total passage number 30 (P30) to minimize genetic drift. Cells were consistently validated for EGFR antigen presence using both antibody and aptamers staining. Consistency in cell morphology was checked prior to each passage. For assays where cells were treated with aptamers, serum concentrations were kept at ∼1% to minimize nuclease exposure.

### Cell viability assay

Cells were seeded at ∼2500 cells/well in a 96-well flat-bottom tissue culture plate 24 h prior to treatment. (In general, “24 h” and “48 h” time points are ± 1-2 hr.) For single dose experiments, cells were treated with aptamers at reported concentrations or with vehicle, and MTT-based cell viability reagent (CellTiter96, Promega, Madison, WI) was added at 48 h and allowed to incubate for ∼2 h before reading. For multidose experiments, cells were treated with 3 µM aptamers or vehicle control in 24 h intervals (media replaced every 48 h), and cell viability reagent was added and read 48 h after the last dose. Absorbance values at 490 nm were obtained and cell viability was reported relative to vehicle. For internalization experiments, endocytosis inhibitor at reported concentrations was added to the well ∼15 min prior to aptamer treatment.

### Cytotoxicity assay

Cells were seeded at ∼10,000 cells/well in a 48-well flat-bottom tissue culture plate 24 h prior to treatment. Cells were treated with aptamers at reported concentrations or with vehicle (DPBS and Mg^2+^) for 48 h and then stained according to the CellEvent™ Caspase-3/7 Green Flow Cytometry Assay Kit (ThermoFisher Scientific, cat: #C10427). Briefly, cells were moved to a 1.5 mL Eppendorf tube, resuspended in Live Cell Imaging Solution (Invitrogen), and stained with CellEvent™ Caspase-3/7 Green Detection Reagent for 30 min and SYTOX™ AADvanced™ Dead Cell Stain for 5 min at 37°C and 5% CO_2_. Cells were immediately analyzed via flow cytometry.

### Western blots

Cells were seeded at ∼8000 cells/cm^2^ in a 12-well flat-bottom tissue culture plate 24 h prior to treatment. Cells were treated with one or two doses of 3 µM aptamers or vehicle for 48 h, one dose of 0.2 µM nimotuzumab for 48 h, or one dose of 0.01 µM Osimertinib for 16 h. For cells treated with two doses of aptamer the second dose was added at 24 h. Cells were washed with ice cold 1X DPBS and lysed in 1X RIPA buffer supplemented with phosphatase and protease inhibitor cocktails on ice for 30 min (with scraping). Cell lysates were either kept on ice and used immediately or stored at −80°C. Prior to gel loading, lysates were quantified using Pierce^TM^ 660nm Protein Assay Reagent (ThermoFisher Scientific cat #22660) and then ∼4 µg of total cell lysate in 1X Laemmli loading buffer were loaded into a 4-12% SurePAGE gel (GeneScript, Piscataway, NJ). PAGE gels were run at 130V for ∼1h in Bis-Tris buffer (GeneScript) and then wet transferred to a 0.2 µm PVDF membrane (BioRad, activated in 100% methanol) at 100V for ∼1 h in 1X transfer buffer containing 20% methanol. Blots were blocked for 1 h at RT in 1X TBST + 3% BSA Fraction V (Gold Biotechnology, Olivette, MO) and then incubated in primary antibody diluted in blocking buffer with overnight rocking at 4°C (phosphorylated and total EGFR, ERK, or AKT) or for 1 h at RT (GAPDH). Blots were washed for 5-10 min in 1X TBST (x3) and incubated in secondary antibodies diluted in blocking buffer for 1 h at RT. Blots were washed again in 1X TBST (x3) and then incubated with SuperSignal^TM^ West Pico PLUS Chemiluminescent substrate (ThermoFisher Scientific) in the dark for 5 min prior to imaging on an Invitrogen^TM^ iBright^TM^ 1500 imaging system. Images were analyzed using FIJI. Blots were stripped using two 5-10 min washes with mild stripping buffer (pH = 2) followed by two 15 min washes with 1X TBST. After stripping, blots were blocked and re-probed as above.

### CFSE proliferation

Approximately 500,000 H1975 or H3255 cells were stained in 1X Carboxy Fluorescein Succinimidyl Ester (CFSE) in DPBS (Biolgend) for 15 min at 37°C and 5% CO_2_, washed with complete medium, and seeded at ∼10,000 cells/well in a 48-well flat-bottom tissue culture plate 24 h prior to experiment. Cells were treated with aptamers at reported concentrations or with vehicle for 24 or 48 h. At reported timepoints, cells were moved to 1.5 mL Eppendorf tubes and analyzed via flow cytometry.

### Cell cycle analysis and Ki67 staining

Cells were seeded at ∼10,000 cells/well in a 48-well flat-bottom tissue culture plate 24 h prior to treatment. Cells were treated with aptamers at reported concentrations or with vehicle for 48 h, moved to a 96 well plate, washed once with DPBS, and then fixed in 70% ice cold EtOH for 2 h. For cell cycle analysis, cells were washed once with DPBS and stained with 100 µg/ml propidium iodide in DPBS supplemented with 50 µg/ml RNase A overnight (∼16 h) at 4°C. For Ki67 staining, cells were washed once with DPBS and stained with APC-labeled anti-Ki67 antibody or isotype control diluted to desired concentration in DPBS supplemented with 0.1% BSA for 1.5 h at room temperature (∼20°C). After staining, cells were washed once with DPBS and analyzed via flow cytometry.

### EGFR cell surface expression

Cells were seeded at 40-50,000 cells/well in a 48-well flat-bottom tissue culture plate 24 h prior to treatment. Cells were treated with aptamers at reported concentrations or with vehicle for 24 or 48 h. At reported time points, cells were washed once with ice cold DPBS, and APC-labeled EGFR or isotype control diluted to desired concentration in DPBS supplemented with 0.1% BSA was incubated for 45 min on ice to prevent internalization. After incubation, antibodies were removed, cells were washed once with DPBS, and cells were lifted off with 1X TrypLE Express (ThermoFisher, Wattham, MA) for <5min at 37°C. TrypLE Express was diluted by the addition of complete medium, and cells were transferred to 1.5 mL Eppendorf tubes and gently centrifuged to pellet (5min, 400x g). Cells were then fixed in 4% paraformaldehyde (PFA) in DPBS in the dark for 5 min at 4°C before being pelleted (5 min, 400x g) and resuspended in DPBS. Cells were kept in the dark at 4°C until analysis with flow cytometry.

### Flow cytometry and analysis

Flow cytometry was performed on an Attune NxT (ThermoFisher Scientific, Waltham, MA) by counting 10,000-30,000 live cell events. For EGFR and Ki67 staining, far-red dyes (APC) were excited using a 637 nm laser, and fluorescence was detected using a standard RL1 filter [670 ± 7 nm]. For CellEvent^TM^ Caspase 3/7 Green Detection Reagent and CFSE staining, cells were excited using a 488 nm laser and detected using standard BL1 filter [530 ± 15 nm]. For SYTOX™ AADvanced™ Dead Cell staining, cells were excited using a 488 nm laser and detected using standard BL3 filter [695 ± 40 nm]. For propidium iodide staining (cell cycle analysis), cells were excited using a 561 nm laser and detected using a standard YL2 filter [620 ± 15 nm]. Flow cytometry data were analyzed and processed using FlowJo Software v10 (Treestar, Ashland, OR). Singlet cells were gated and analyzed for mean fluorescence intensity (MFI, antibody and CFSE staining) or percent positivity (Caspase 3/7 and Sytox staining). For cell cycle analysis, a Watson (Pragmatic) univariate model with a G1/G2 peak coefficient of variation (CV) = n = 9 was used to differentiate propidium iodide staining.

### Subcutaneous xenograft of NSCLC cells

To develop cell-line-derived subcutaneous tumor models^98^, H1975 cells were grown as described above in 15 cm dishes to ∼80% confluence. Cells were collected, washed twice with cold DPBS, and resuspended in ice cold DPBS plus 50% extracellular matrix (ECM) gel (Sigma Aldrich, cat: #E6909) at ∼5 million cells per 100 µL. Cells were kept on ice during injections to prevent solidification of ECM gel. To develop xenografts, BALB/c nude mice (Charles River, Wilmington, MA) were briefly anesthetized with isoflurane in an induction chamber (4%, 250 mL/s). Using a 28G 1/2” 1 cc insulin syringe/needle, 5×10^6^ cells (100 µL) were subcutaneously injected into the flank of each mouse. Cells were allowed to engraft for one week and then mice were monitored daily until all tumors could be palpated and measured with digital calipers (minimum size ∼4 mm^3^).

### *In vivo* tumor burden

Aptamers for *in vivo* experiments were folded at 3 µM in DPBS without MgCl_2_. On day 0, nude mice harboring barely palpable subcutaneous tumors with sizes averaging 13.3 ± 4.8 mm^3^ were briefly anesthetized with isoflurane in an induction chamber (4%, 250 mL/s). Using a 28G 1/2″ 1cc insulin syringe/needle, 50 µl (∼150 pmol; day 0, 2, 4) or 65 µl (∼200 pmol; day 6, 7) of aptamer solution or vehicle (DPBS only) was injected intratumorally. Tumor growth and mouse weight were monitored every 1-3 days until at least one mouse reached the humane endpoint (a diameter >20 mm or volume >2000 mm^3^). No mice experienced significant weight changes (data not shown). Tumor diameters for width (W) and length (L) were measured and raw tumor volumes (mm^3^) were calculated using the following equation: V = 0.5W * W * L. Reported relative tumor volumes were normalized to tumor size at day 0. At the endpoint (day 17) mice were humanely sacrificed and imaged in prone and right recumbent positions. Tumors were resected and *ex vivo* endpoint tumor volumes and weights were measured. Tumors were placed in 1.5 mL Eppendorf tubes, snap frozen in liquid nitrogen, and stored at −80°C for downstream analyses.

### Animal ethics statement

All animal procedures were conducted according to NIH guidelines for the care and use of laboratory animals and were approved by the University of Missouri Institutional Animal Care and Use Committee.

### Bulk transcriptomics sample preparation

H3255, H1975, H820, and A549 cells were seeded at ∼10,000 cells/well in a 48-well flat-bottom tissue culture plate 24 h prior to treatment. Cells were treated with 3 µM *EGFRapt* (MinE07), control apt (C36), or vehicle (DPBS and Mg^2+^) for 24 h, after which the cells were washed twice with DPBS and total RNA was extracted using the Monarch® Total RNA Miniprep Kit (NEB, Ipswich, MA). Total RNA was stored at −80°C until sequencing. Illumina NGS was performed on total RNA from three biologically independent experiments by the University of Missouri DNA Genomics Technology Core using NovaSeq technology to provide 100 base paired end (PE) reads.

### Bulk transcriptomics differential expressions analysis

The bulk transcriptomic FASTQ files for each of the four cell lines (H3255, H1975, H820 and A549), for three conditions (EGFRapt, Vehicle, Control), with three replicates each underwent quality check using FastQC (v0.12)^99^. Consequently, the individual results were aggregated by MultiQC (v0.4) to generate a comprehensive report of Sequence Quality Histograms, Per Sequence Quality Scores, Per Sequence GC Content, Per Base N Content, and Adapter Content. To process bulk RNA sequencing data, Python language v3.7 was used to integrate command line tools to run together as an automated pipeline. Sequences were trimmed using Trim Galore^100^ and aligned (mapped) via Hisat2 (v2.2.1)^101^ with reference genome file “GR38_latest_genomic.fna” downloaded from (https://ftp.ensembl.org/pub/release-110/fasta/homo_sapiens/dna/). The results of alignment were sorted using samtools (v1.9), followed by transcriptome assembly using Cufflinks (v2.21)^102^. The annotation file used for this step “GRCh38_latest_genomic.gff” downloaded from (https://ftp.ensembl.org/pub/release-111/gtf/homo_sapiens/).

To assess the quality of biological replicates for each condition, Principal Component Analysis (PCA) was performed, and Pearson Correlation Coefficient based clustering matrix was generated using the FPKM (Fragments Per Kilobase of transcript per Million mapped reads) values produced by the Cufflinks suite. The thresholds, |log2FC| >=0.5 and q value <=0.05 were used to identify differentially expressed genes (DEGs), using Cuffdiff (v2.2.1)^102^. Upset plots were generated to display the number of intersecting genes between cell lines for main comparison (*EGFRapt versus control*), as well as number of genes within each condition comparisons, for each cell line. Additionally, hierarchical clustering in combination with heatmap was performed, showing forty-three DEGs associated with cell cycle pathways and their respective log2FC using ComplexHeatmap (v2.18) R package.

For DEGs identified in each cell line, within each set of comparison *EGFR apt* versus Vehicle or Control, functional enrichment analysis was conducted using GProfiler2 (v0.2.2)^102^ R package. GProfiler uses a gene list to perform functional enrichment analysis, retrieving results from sources, Gene Ontology (GO), WikiPathways and Reactome Pathways. Simultaneously, Gene Set Enrichment Analysis (GSEA) was carried out using fgsea (v1.28.0)^103^ R package, which evaluates the overrepresentation of genes based on a pre-ranked gene list, where the ranking of genes is determined on the bases of its correlation to a particular phenotype, in this case condition within a cell line.

The data of differentially expressed genes (DEGs) can be accessed and analyzed interactively through the integrated 3DOmics tool in HumanKB within KBCommons^104,105^ framework (https://kbcommons.org/system/browse/diff_exp/homoSapiens). The tool allows comparison of various experimental conditions together, visualization of upsets, venn diagram and volcano plots. In addition, it is also possible to perform functional enrichment analysis using GProfiler creating a one stop, integrated solution for bioinformatics differential expression analysis.

---

**Figure S1.**
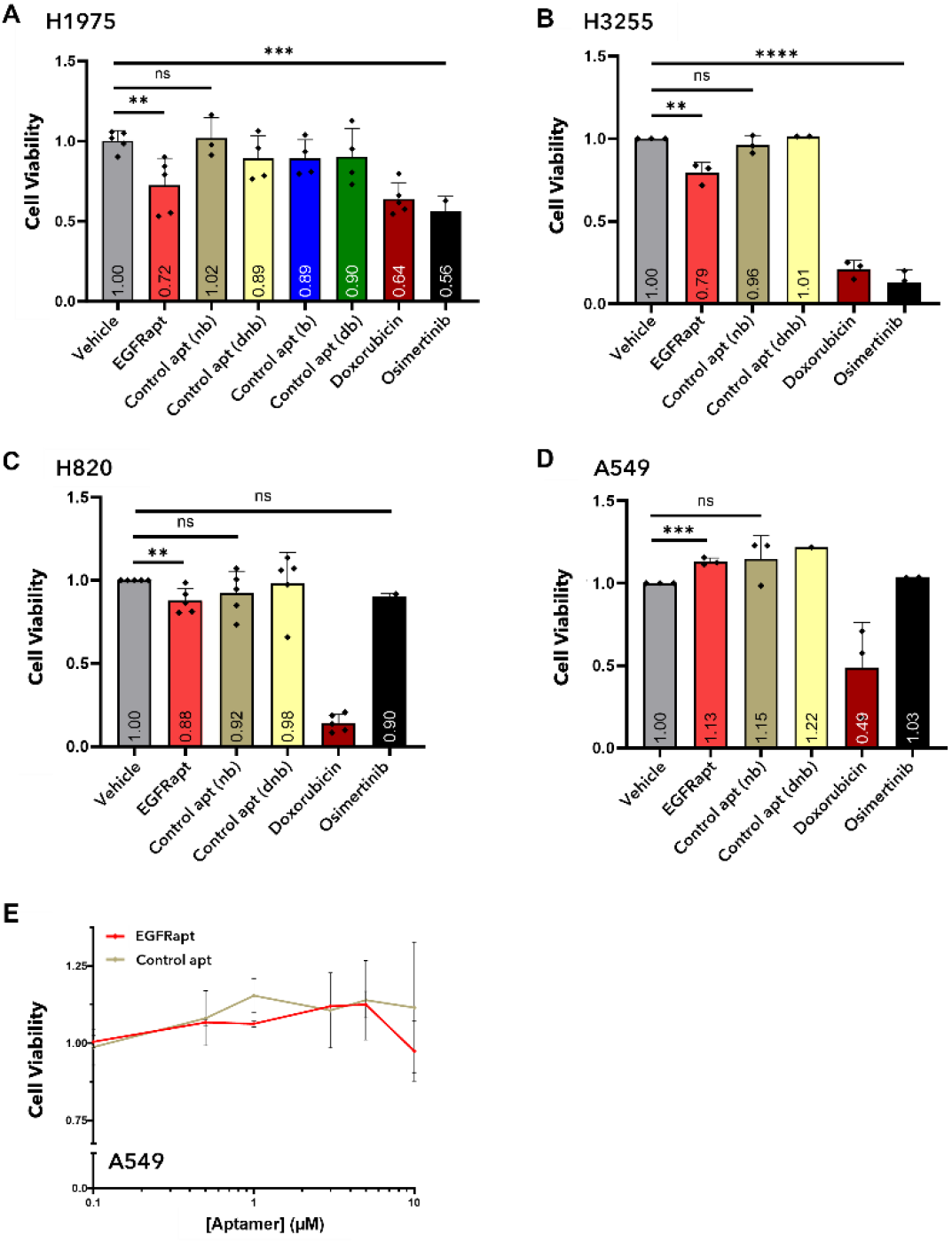
Effect of *EGFRapt* on cell viability of NSCLC cell lines. (**A-D**) All cells were subject to cell viability assay as in Figure 1B. H1975 (A), H3255 (B), H820 (C), or A549 (D) were treated with 3 µM *EGFRapt* (red bar), 2’FY RNA non-binding (nb, tan bar; extended version of C36), DNA non-binding (dnb, yellow bar; scrambled DW4), 2’FY RNA binding (b, blue bar; extended version of Waz), or DNA binding (db, green bar; minimized CLN3) control apts, 5 µM Doxorubicin (dark red bar), 0.5 µM Osimertinib (black bar), or vehicle (grey bar) for 48 h. Cell viability was determined using MTS reagent and plotted relative to viability of vehicle-treated cells. *EGFRapt* decreased cell viability in H1975 and H3255 cells but not in H820 or A549 cells when compared to non-binding control apts. (**E**) A549 cells were subject to cell viability assay as in Figure 1C. *EGFRapt* (red line) had no effect on cell viability in A549 cells at doses up to 10 µM compared to non-binding control apt (tan line). For A-E, relative cell viability (normalized to vehicle) is reported on the *y*-axis and plotted values represent mean ± SD for n=2-5 independent experiments, each with 2-4 technical replicates. Statistical analysis was preformed using a two tailed t-test (*p < 0.05).

**Figure S2.**
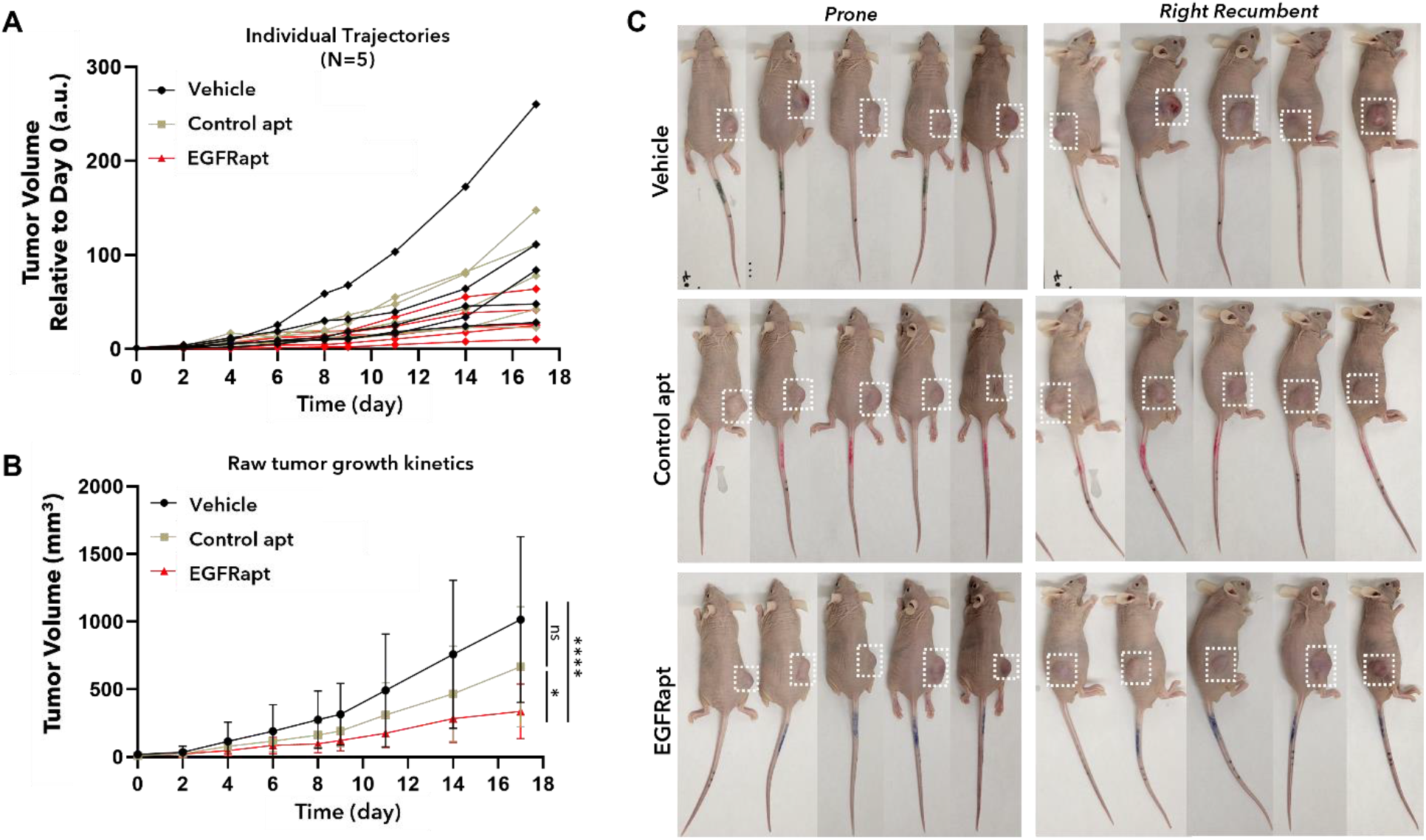
*EGFRapt* decreases tumor burden in nude mice bearing NSCLC cell line derived xenografts. (**A-C**) Mice were treated as in Figure 2. Tumor bearing mice were treated with five intratumoral doses of *EGFRapt* (red line), control apt (tan line), or vehicle (black line) and tumor burden was tracked until a humane endpoint. (**A**) Raw tumor volume (mm^3^) is plotted on the *y*-axis and plotted values represent mean ± SD for n=5 mice from each treatment group. (**B**) Individual tumor trajectories for n=5 mice from each treatment group are shown where tumor volume (mm^3^) relative to day 0 is plotted on the *y*-axis. (**C**) Mice from each treatment group were imaged in the prone (left) and right recombinant (right) positions at the endpoint. Tumor growth kinetics were significantly decreased in mice treated with *EGFRapt* compared to control apt or vehicle (PBS). Statistical analysis was preformed using a simple linear regression model (*p < 0.0332, ****p < 0.0001)

**Figure S3.**
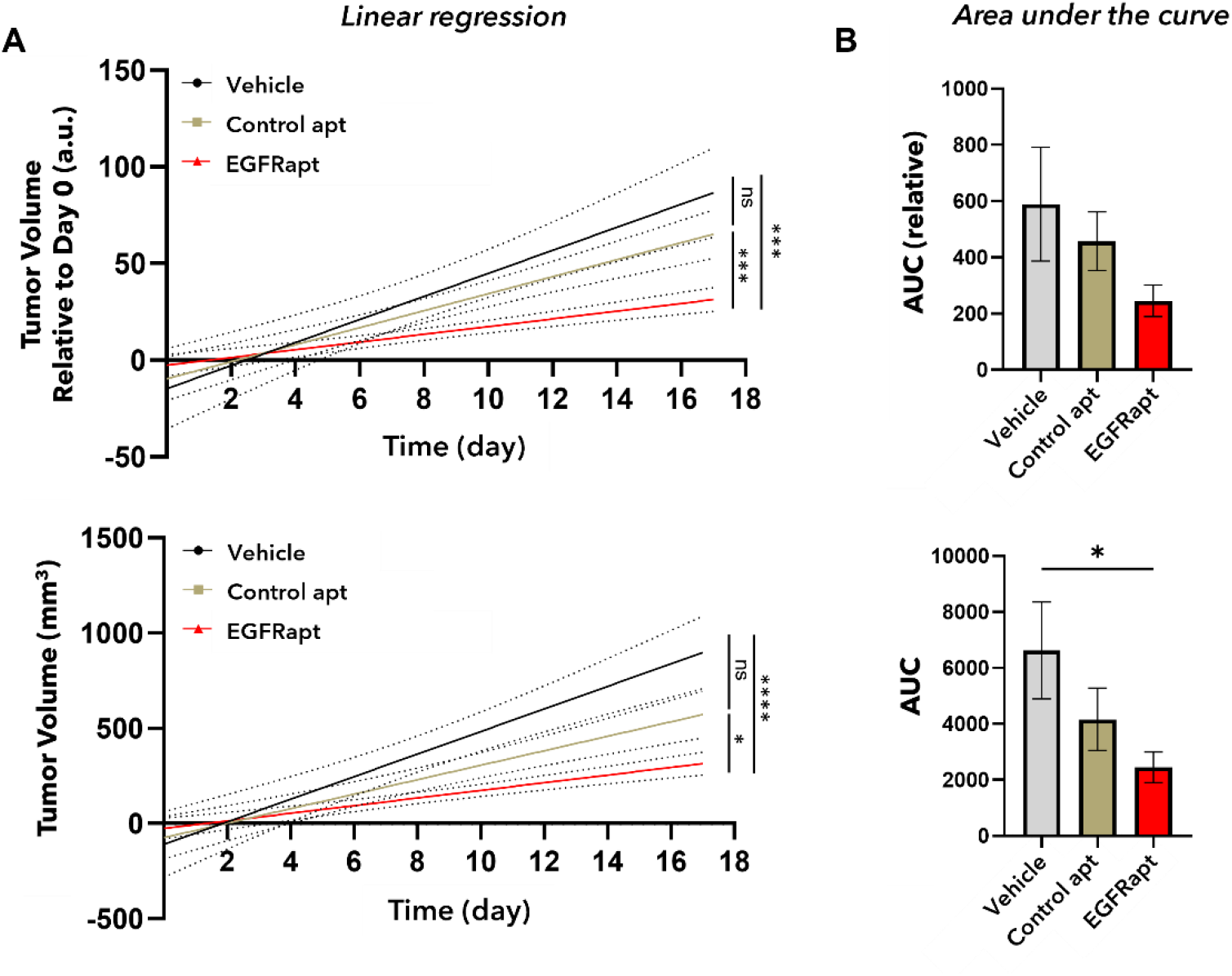
Statistics for normalized and raw tumor growth data. (**A-B**) Statistical analyses of tumor burden upon treatment with *EGFRapt*, control apt, or vehicle (Figure 2 and S2; n=5 mice for each treatment group). Simple linear regression (A) and a two tailed t-test of area under the curve (AUC; B) are shown. Using a linear regression model, treatment with *EGFRapt* (red line/bar) was significant for normalized (*top*; relative to day 0) and raw (*bottom*) data when compared to vehicle (grey line/bar) or control apt (tan line/bar; *p < 0.0332, ***p < 0.0002, ****p < 0.0001). Using an AUC model, treatment with *EGFRapt* was significant only for raw data when compared to vehicle (*p < 0.05).

**Figure S4.**
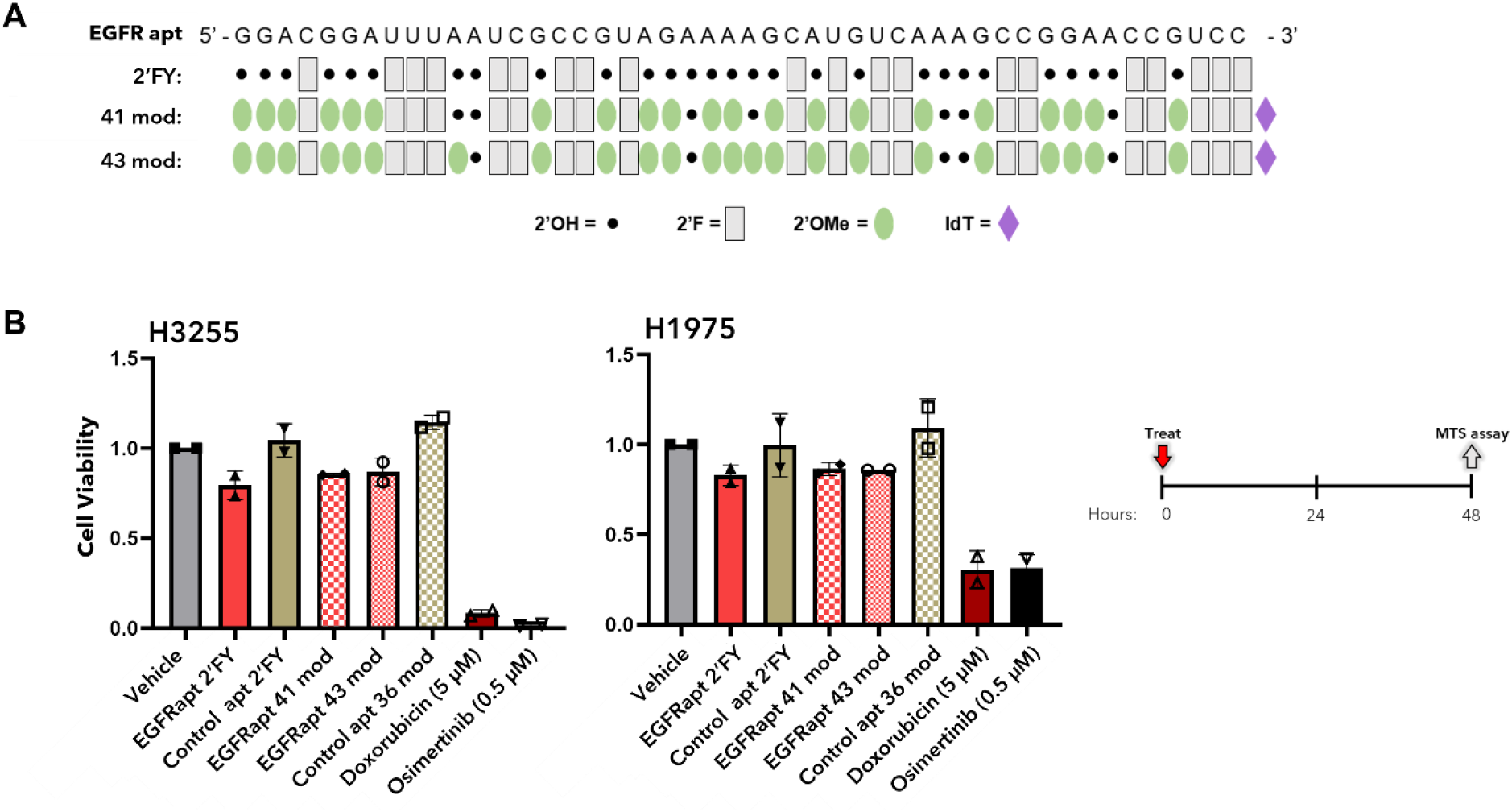
Highly modified versions of *EGFRapt* similarly decrease cell viability in NSCLC cell lines harboring L858R mutations. (**A**) Schematic of original (2’FY) and highly modified versions (36 and 38 mod; patent US10190121B2) of *EGFRapt*. Black dots represent canonical 2’ hydroxy (OH), grey squares represent 2’fluoro (F) modifications, green ovals represent 2’ O-methyl (OMe) modifications, and purple diamonds represent 3’ inverted dT (IdT) modifications. Modified *EGFRapt*s tested were selected from patent US10190121B2 (36.01 and 38.01) [51]. (**B**) H3255 (*left*) and H1975 (*right*) cells were treated with 3 µM original *EGFRapt* (solid red bar) or control apt (solid tan bar) or highly modified versions of *EGFRapt* (36 and 38 mod; red/white bars) or control apt (tan/white bar) for 48 h and cell viability was determined using MTS reagent (*assay schematic*). Relative cell viability (normalized to vehicle) is reported on the *y*-axis. Plotted values represent mean ± SD for n=2 independent experiments, each with 2-4 technical replicates. In both cell lines, highly modified versions of *EGFRapt* that contain 2’OMeR modifications and a 3’ IdT decreased cell viability similar to the original *EGFRapt*.

**Figure S5.**
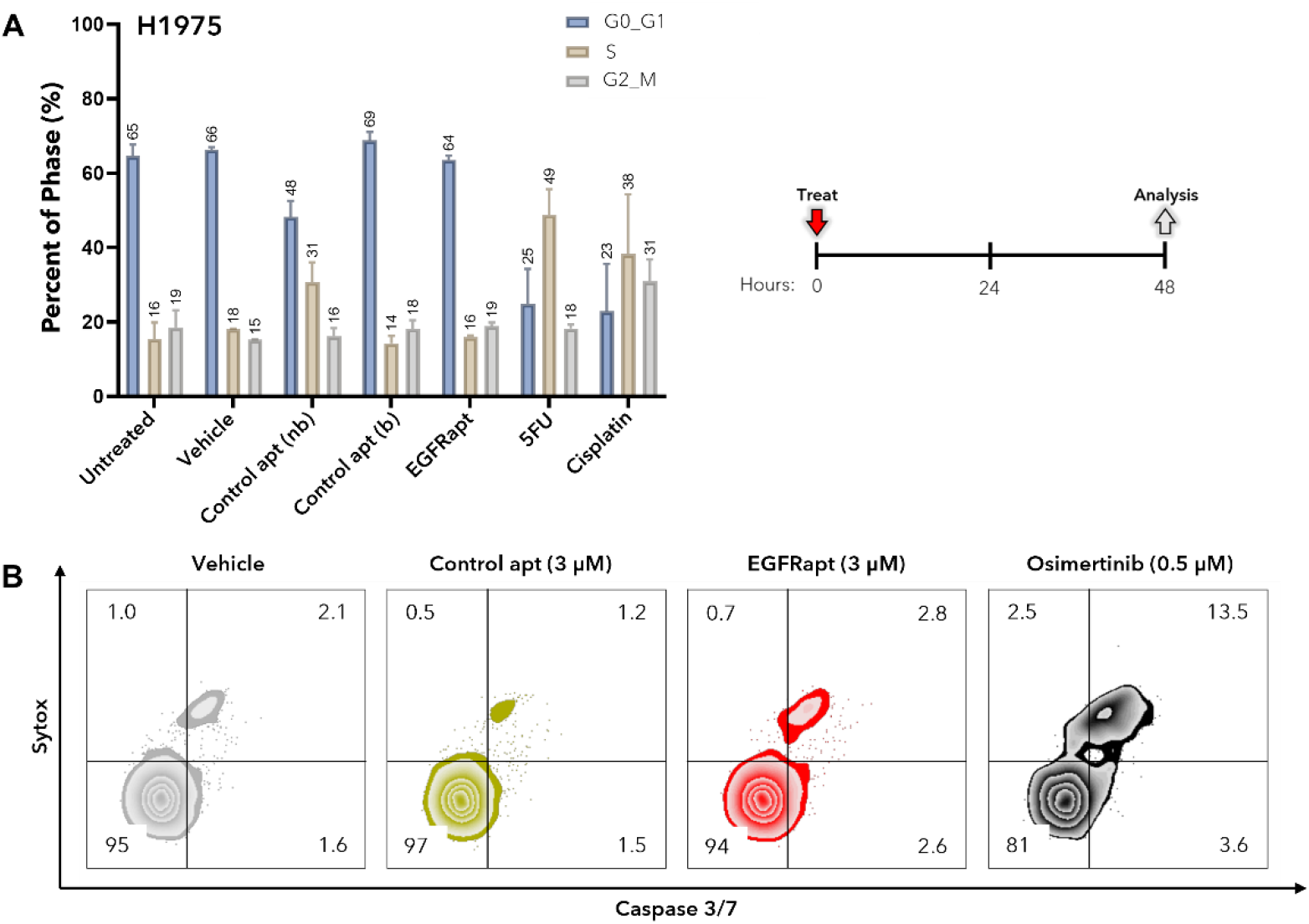
*EGFRapt* does not induce cell cycle arrest and is not cytotoxic in H1975. (**A**) H1975 cells were treated with reported concentrations of *EGFRapt*, control apt, or chemotherapeutic for 48 h and then fixed and stained with propidium iodide. Cell cycle determination was analyzed by flow cytometry and using the Watson pragmatic method. The percentage of cells in the G0-G1 (blue bar), S (sand bar), or G2-M (grey bar) phase of cell cycle is plotted on the *y*-axis. Plotted values represent mean ± SD for n=2-3 independent experiments. H1975 cells treated with *EGFRapt* did not alter cell cycle phase transition compared to binding (b) control apt or vehicle. (**B**) H1975 cells were treated as in A but were live stained with apoptosis marker (CellEvent^TM^ Caspase 3/7 reagent) or cell impermeable necrosis and late apoptosis marker (Sytox^TM^). Cell staining was analyzed by flow cytometry. Representative contour plots in which percent of cells unstained (Q3) or stained with Caspase 3/7 reagent only (Q4), Sytox^TM^ only (Q2), or both reagents (Q1) from one of n=2 independent experiments is shown. H1975 cells treated with *EGFRapt* had no effect on Caspase 3/7 reagent or Sytox^TM^ staining when compared to non-binding control apt or vehicle.

**Figure S6.**
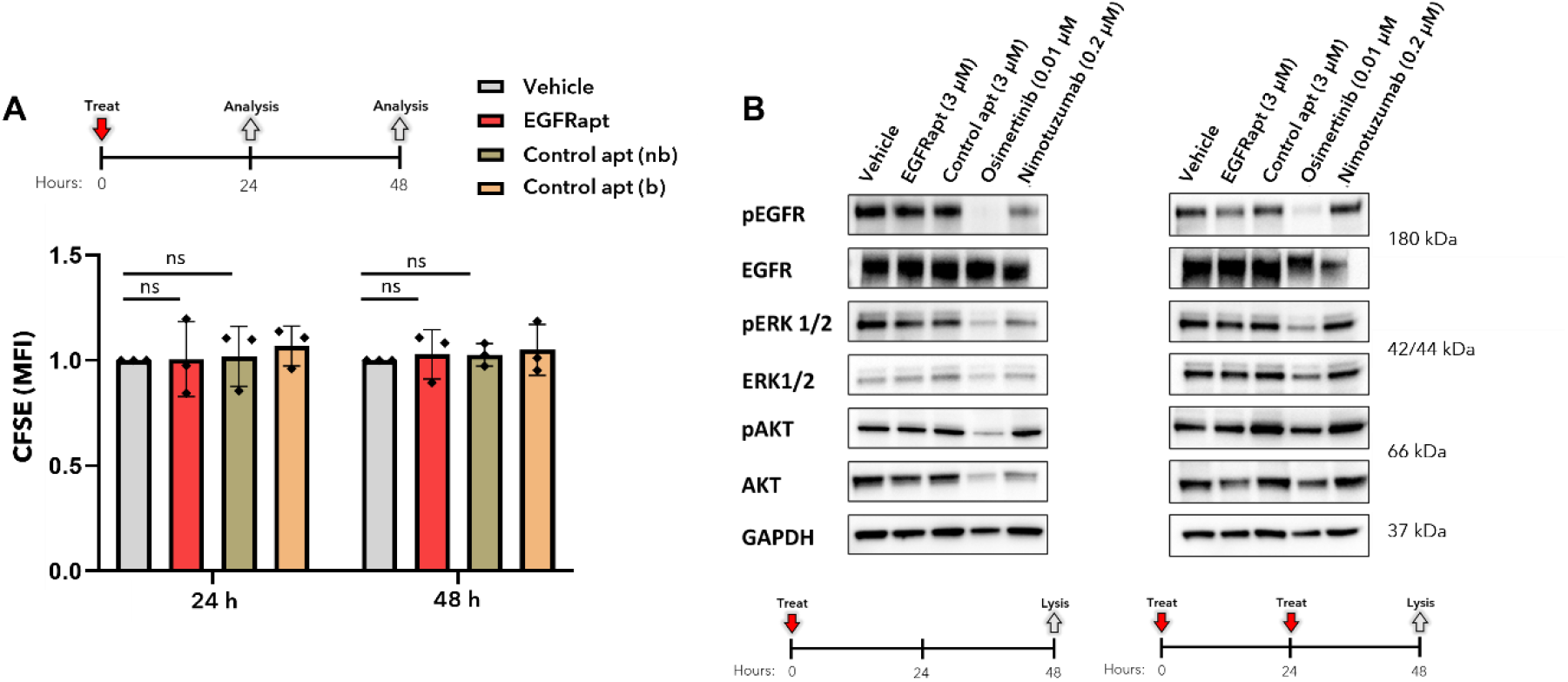
*EGFRapt* does not alter canonical EGFR signaling or proliferation in H3255. (**A**) H3255 cells were prelabeled with Carboxy Fluorescein Succinimidyl Ester (CFSE) and treated with 3 µM *EGFRapt* (red bar), binding (b; orange bar) or non-binding (nb; tan bar) control apts, or vehicle (grey bar) for 24 or 48 h and then analyzed by flow cytometry (*assay schematic*). The mean fluorescence intensity (MFI) of CFSE staining is plotted on the *y-*axis. H3255 cells treated with *EGFRapt* exhibited no change in CFSE staining compared to control apts, suggesting no change in proliferation rates. (**B**) Total cell lysates from H3255 cells treated with the indicated doses of one (*left*) or two (*right*) doses of *EGFRapt*, control apt or vehicle for 48 h, one dose of Nimotuzumab for 48 h, or one dose of Osimertinib for 16 h were probed for total and phosphorylated EGFR, ERK1/2, AKT, or loading control, GAPDH. Quantification of blot intensity (data not shown) revealed little to no differences in proteins involved in EGFR kinase dependent signaling upon treatment with *EGFRapt* compared to control apt or vehicle.

**Figure S7.**
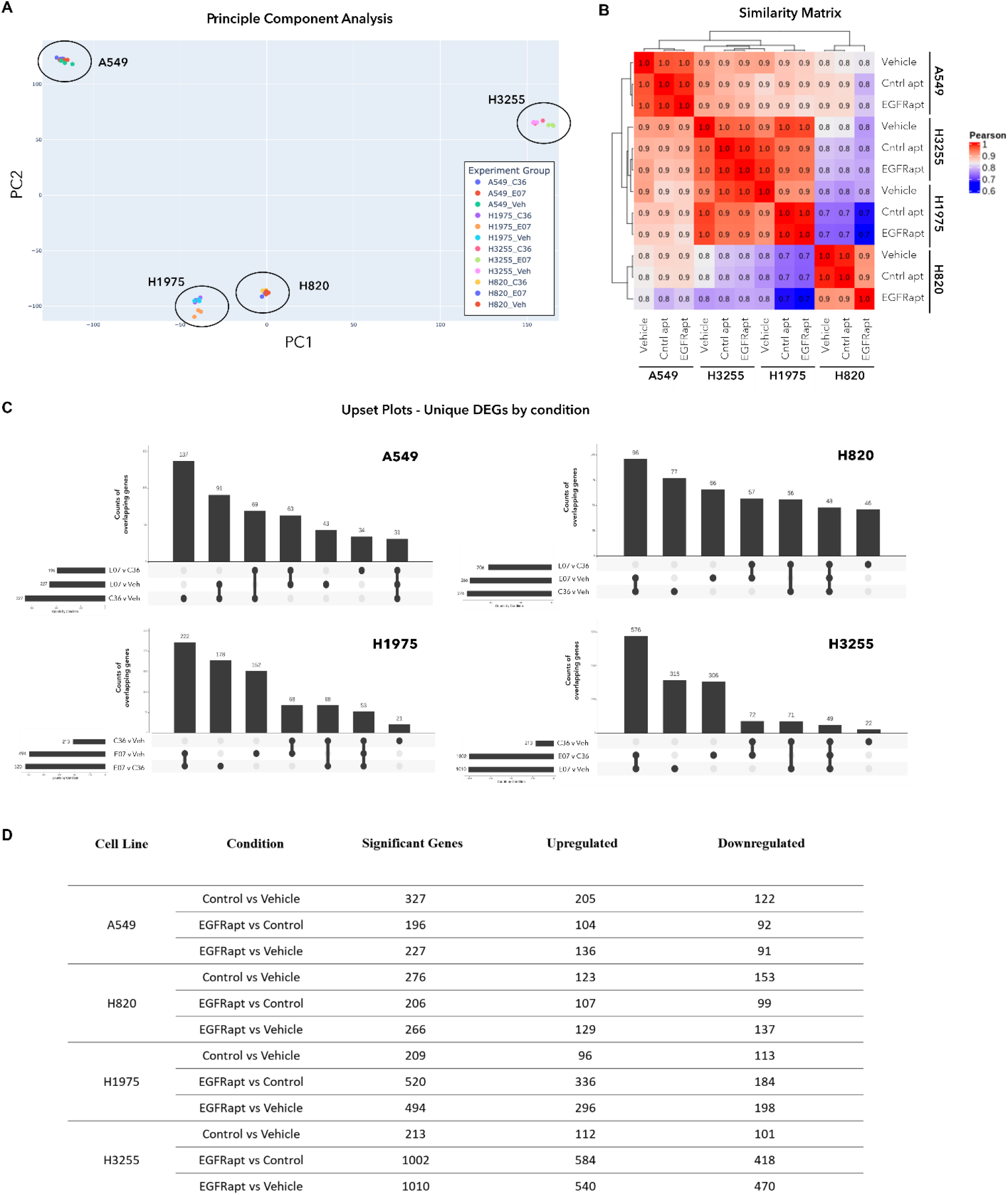
Differentially expressed genes (DEGs) in NSCLC cell lines. (**A**) Linear dimensionality reduction using principal component analysis (PCA) and (**B**) similarity matrix heatmap (red = high correlation; blue = low/moderate correlation) for all cell lines and all treatments. (**C**) Upset plots showing number of unique differentially expressed genes (DEGs) by condition for each cell line. (**D**) Number of upregulated and downregulated DEGs by condition for each cell line. For A and C, E07 = EGFRapt; C36 = control apt; Veh = vehicle. The high-level analyses in A-D are suggestive of cell line-based clustering rather than treatment-based clustering.

**Figure S8.**
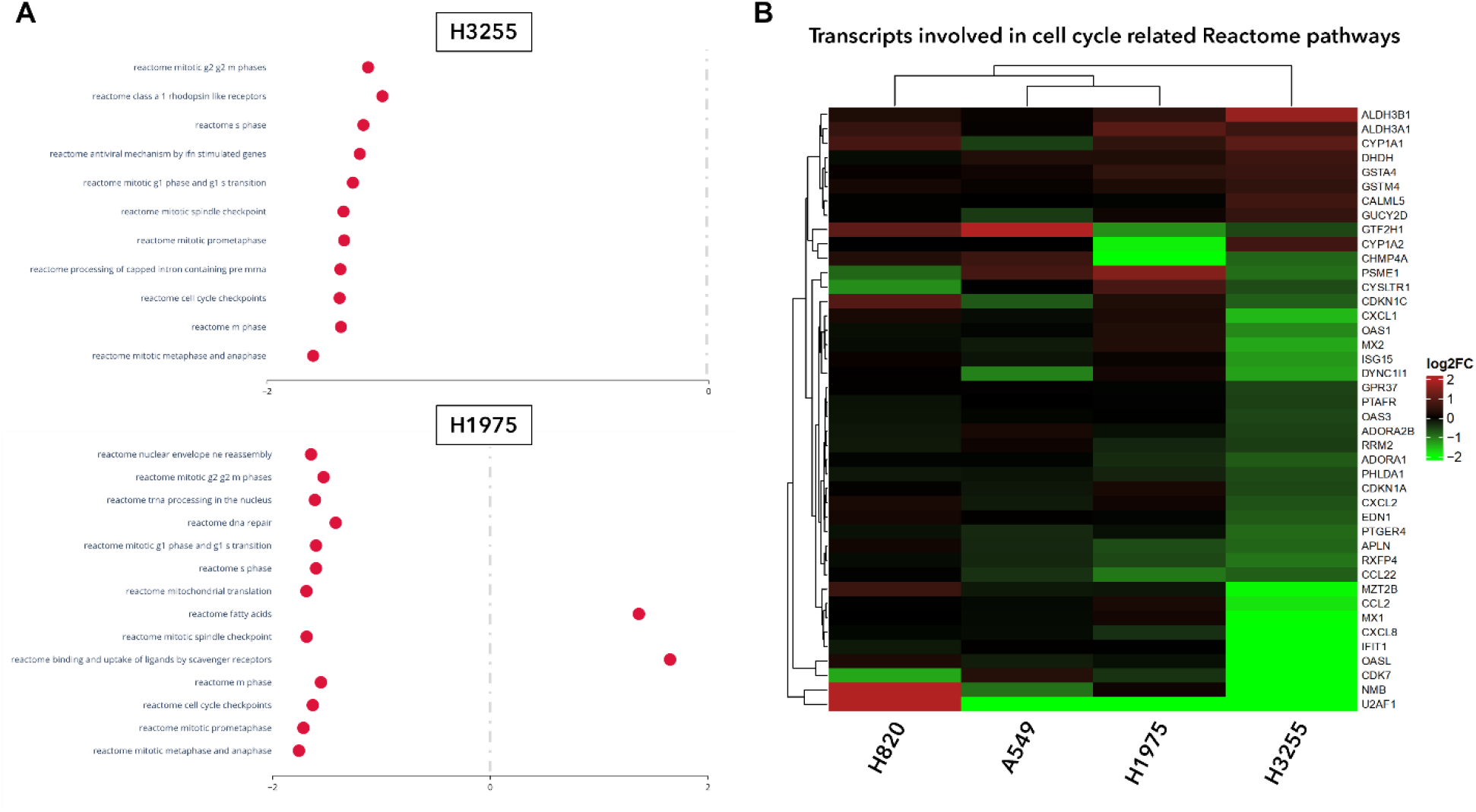
Differentially expressed genes (DEGs) in NSCLC cell lines. (**A**) Normalized enrichment scores (NES) for significantly enriched Reactome and KEGG pathways (GSEA) noted in Figure 5D. (**B**) Heatmap showing relative expression of genes involved in cell cycle related Reactome pathways noted in A and Figure 5D (red = upregulated; green = downregulated; black = no change). NES for cell cycle related Reactome terms in both H3255 and H1975 treated with EGFRapt compared to control apt were negative; however, cell cycle related gene expression was remarkably more downregulated in H3255 compared to H1975.

**Figure S9.**
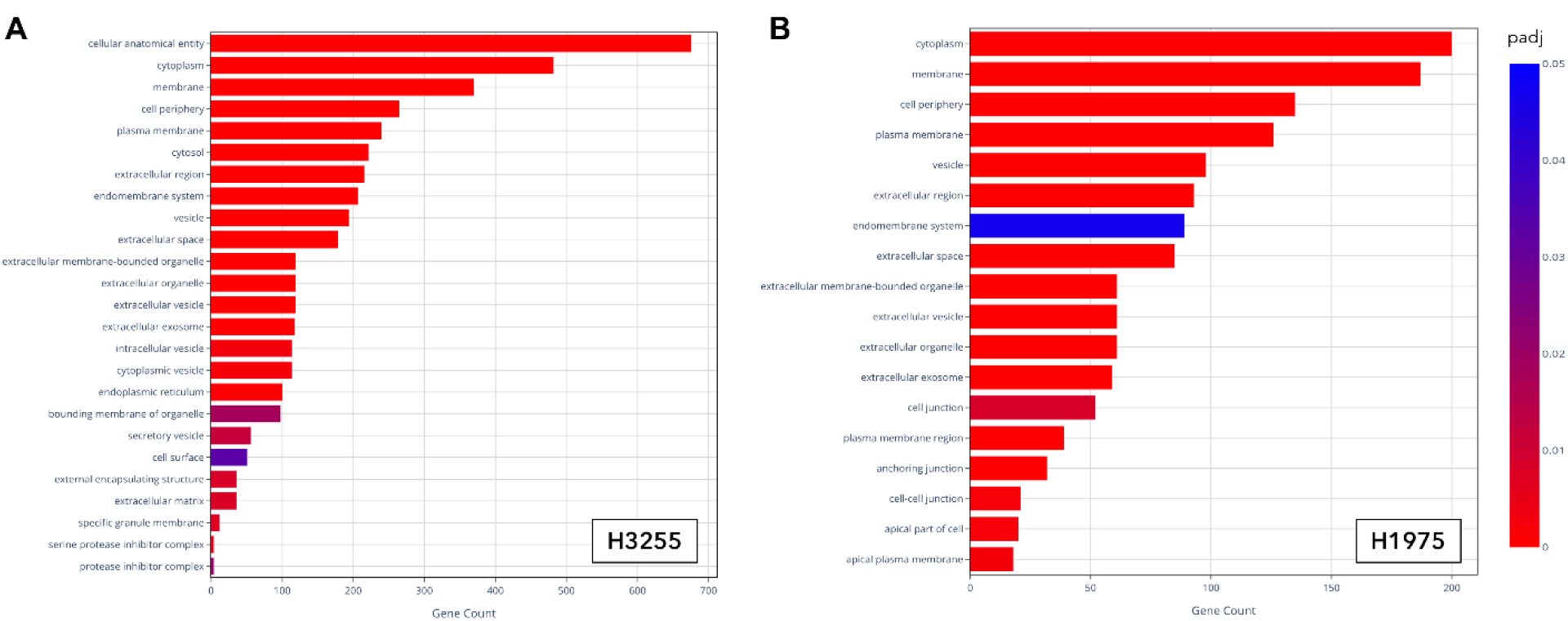
Cytoplasm, membrane, and vesicles are among GO:CC terms enriched in both H3255 and H1975. (**A-B**) Enriched gene ontology terms associated with cellular component (GO:CC) in H3255 (A) and H1975 (B). Terms related to cytoplasm, cell periphery, plasma membrane, and vesicles are significantly enriched in both cell lines treated with EGFRapt compared to control apt.

**Figure S10.**
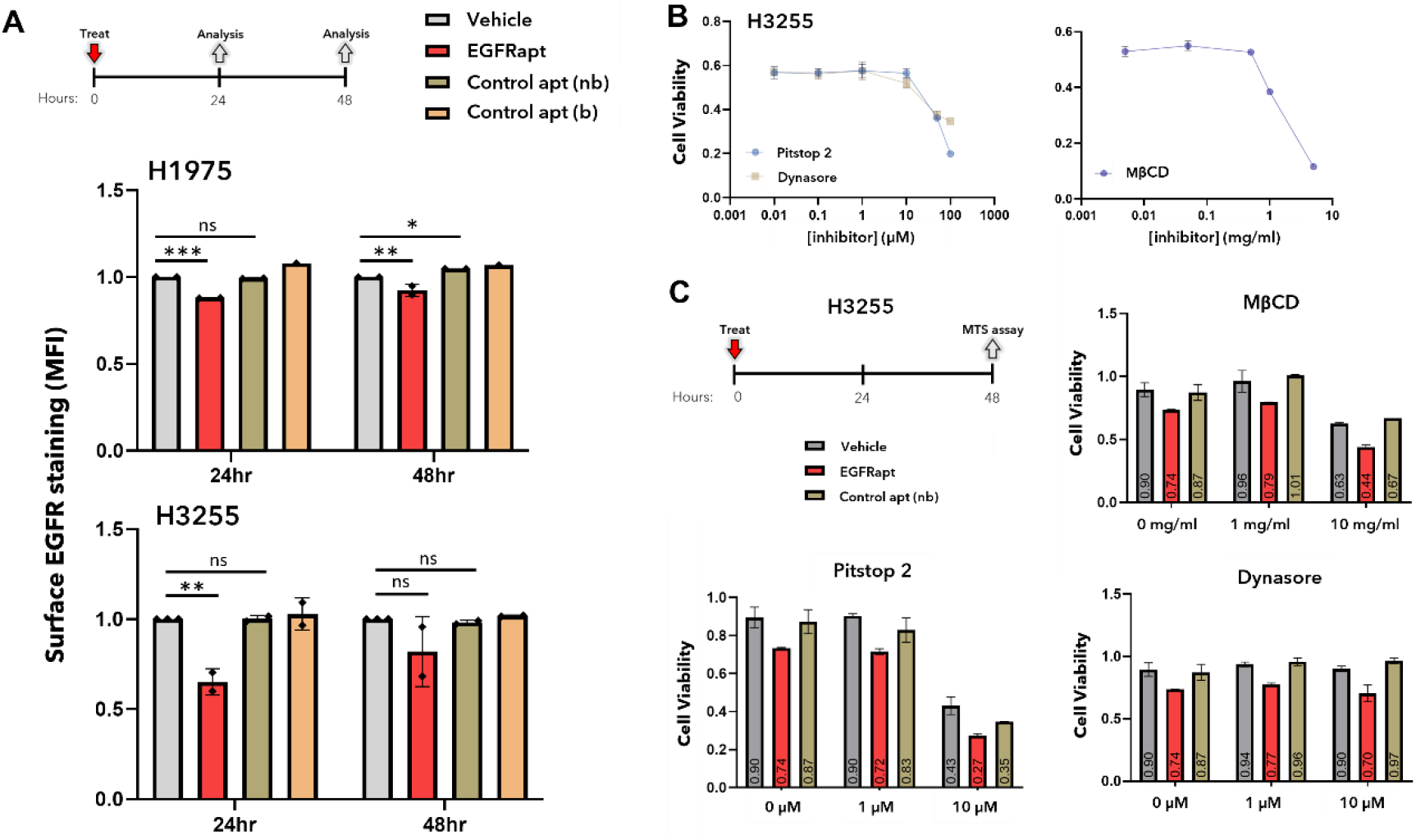
*EGFRapt* decreases cell surface labeling of EGFR and anti-cancer effect is not affected by endocytosis inhibitors. (**A**). H1975 (*top*) or H3255 (*bottom*) were treated with treated with 3 µM *EGFRapt* (red bar), binding (orange bar) or non-binding (tan bar) control apts, or vehicle (grey bar) for 24 or 48 h and then stained with a non-competitive, APC labeled EGFR antibody on ice to prevent internalization (*assay schematic*). Antibody staining was analyzed by flow cytometry. MFI of EGFR antibody (cell surface staining) is reported on the *y*-axis and plotted values represent mean ± SD for n=2 independent experiments (n=1 for binding control apt in H1975). In both cell lines, *EGFRapt* decreased cell surface staining at 24 h and to a lesser extent at 48 h. Statistical analysis was performed using a two-tailed t-test (**p < 0.01, ***p < 0.001). (**B**) H3255 cells were treated with various doses of endocytosis inhibitors—Pitstop 2 (blue line, *left*), Dynasore (sand line, *left*), or methyl-beta-cyclodextrin (MβCD; blue line, *right*) for 48 hr and cell viability was determined by MTS assay. All endocytosis inhibitors negatively impacted cell viability at higher concentrations. (**C**) H3255 cells were treated with reported doses of endocytosis inhibitors in the presence of *EGFRapt* (red bar), non-binding control apt (tan bar), or vehicle (grey bar) for 48 h. Cell viability was determined by MTS assay. In each case, the red bar is below the grey and tan bars, indicating that none of the endocytosis inhibitors altered the anti-cancer effect of *EGFRapt*, even at concentrations that negatively impacted cell viability. For B-C, relative cell viability (raw data in B, normalized to vehicle in C) is reported on the *y*-axis and plotted values represent mean ± SD for n=2 independent experiments, each with 2-4 technical replicates.

**Figure S11.**
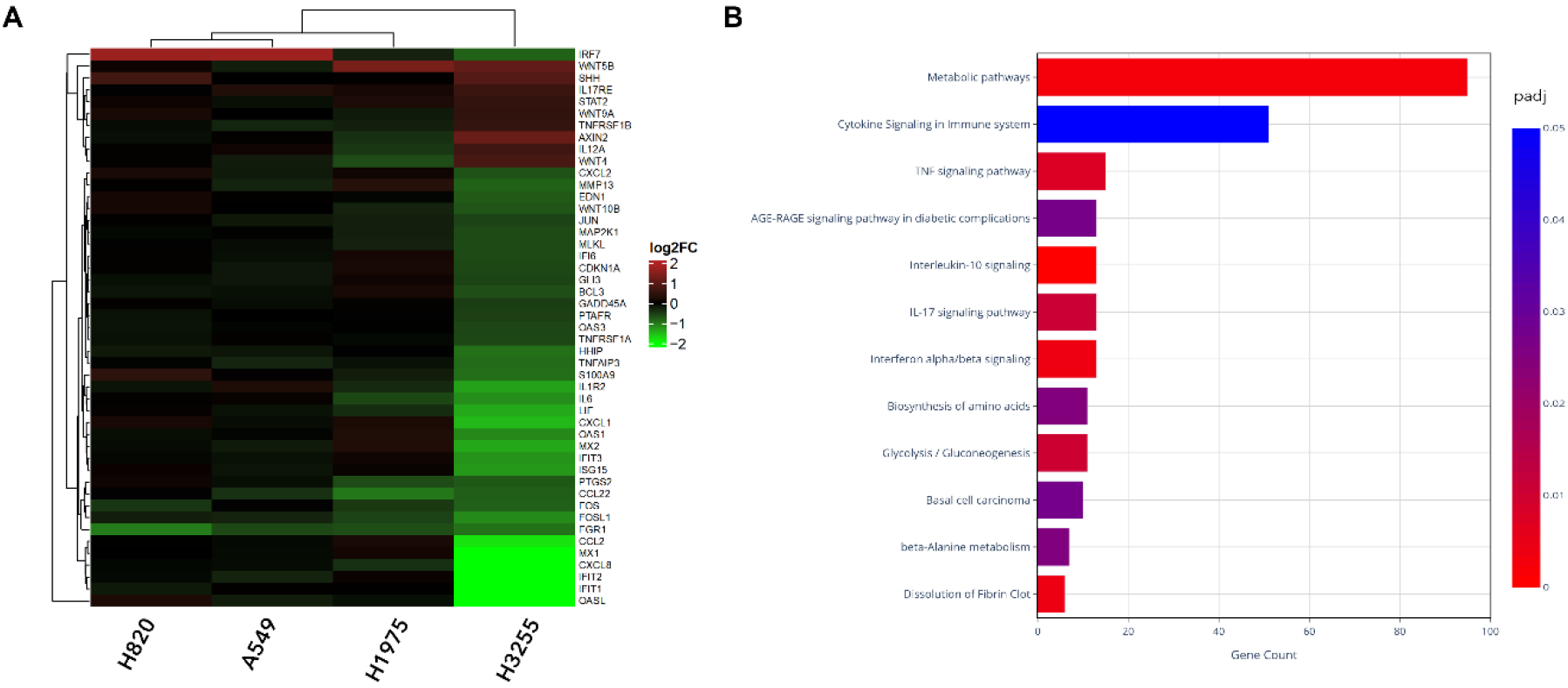
Down regulation of genes involved in interferon and cytokine signaling in H3255. (**A**) Heatmap showing relative expression of genes involved in the immune response in all cell lines (red = upregulated; green = downregulated; black = no change) (**B**) Significantly enriched immune related gene ontology terms associated with biological processes (GO:BP) in H3255. H3255 cells treated with EGFRapt compared to control apt showed down regulation of many genes involved in interferon and cytokine signaling, revealing possible mediators of cytotoxic response observed in this cell line.

**Table S1.**
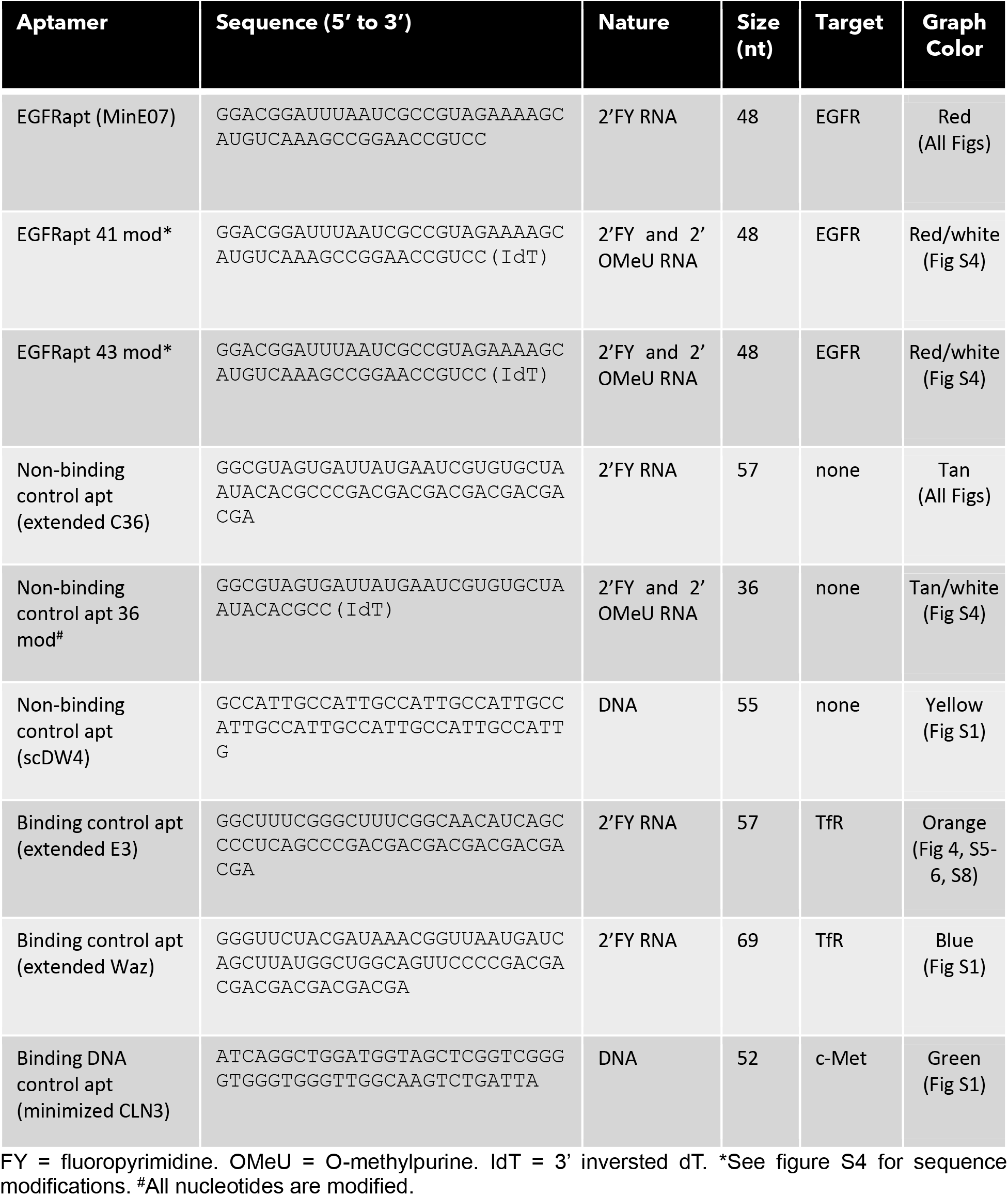
Aptamer sequences.

**Table S2.**
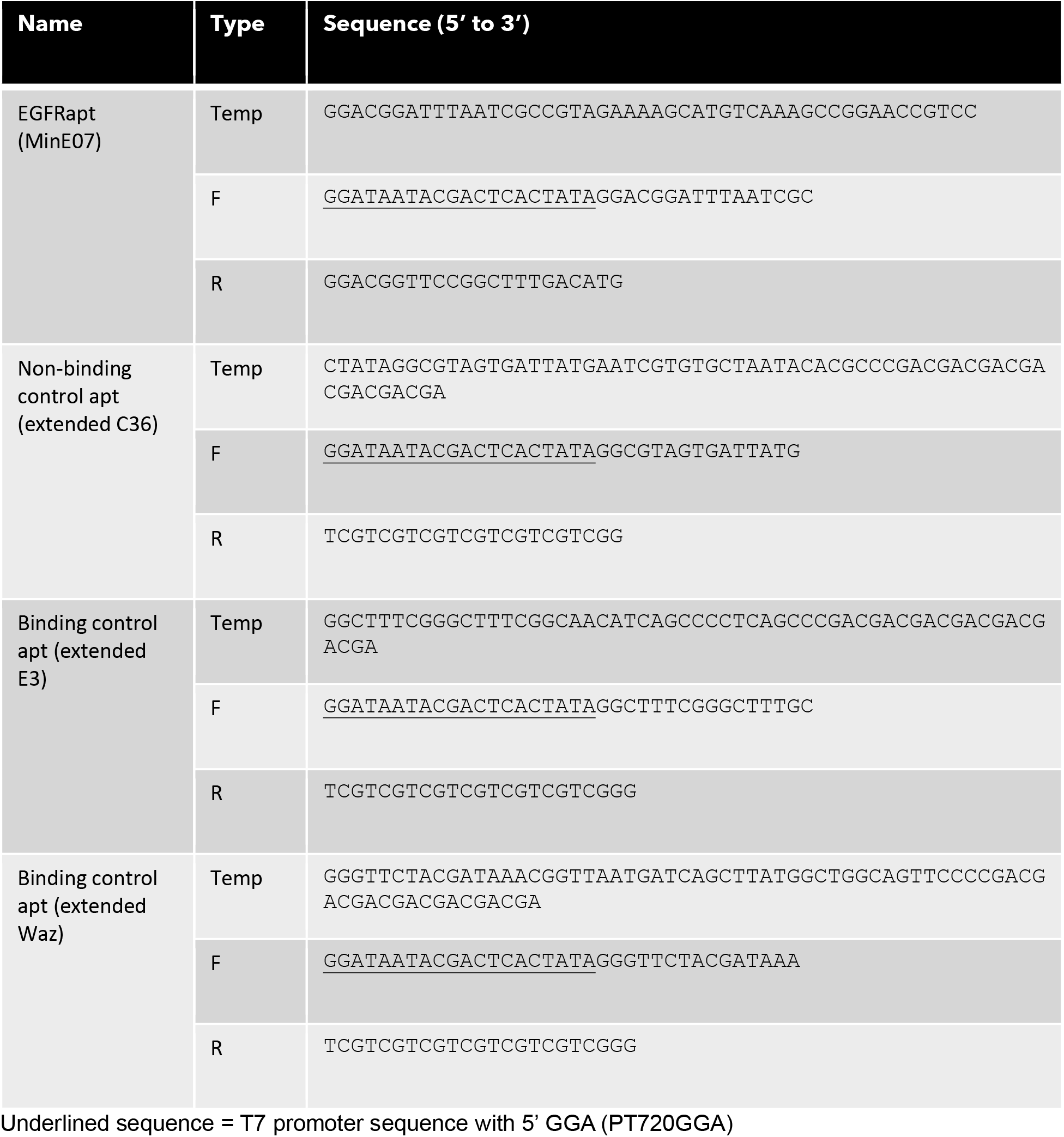
Template and primer sequences for *in vitro* transcription of 2’FY RNA aptamers.

